# Adenita: Interactive 3D modeling and visualization of DNA Nanostructures

**DOI:** 10.1101/849976

**Authors:** Elisa de Llano, Haichao Miao, Yasaman Ahmadi, Amanda J. Wilson, Morgan Beeby, Ivan Viola, Ivan Barisic

## Abstract

We present Adenita, a novel software tool for the design of DNA nanostructures in a user-friendly integrated environment for molecular modeling. Adenita is capable of handling large DNA origami structures, re-use them as building blocks of new designs and provide on demand feedback, thus overcoming effectively some of the limitations of existing tools. Additionally, it integrates all major established approaches to DNA nanostructure design (DNA origami, wireframe nanostructures and DNA tiles) and allows to combine them. We showcase Adenita by re-using a large nanorod designed with Cadnano [1] to create a new nanostructure through user interactions that employ different editors to modify the original nanorod.

## 1 Introduction

DNA origami [2] is currently one of the most popular techniques for the design of DNA nanostructures. It employs a long DNA single-strand, or “scaffold”, that is folded into a predefined nanostructure with the help of several shorter single-strands, or “staples”, which bind to the scaffold at specific positions. Although DNA origami was created to build solid 2D faces, it was soon extended to 3D [3] and to wireframe nanostructures [4, 5]. DNA origami has been successfully applied to create measurement devices [6], enzymatic cascades [7], DNA nanopores [8], biosensing devices [9] and drug delivery vessels [10, 11], amongst others.

The construction of DNA nanostructures usually involves the routing of a long scaffold (approximately 8,000 nucleotides), the placement of the staples and the determination of their sequences. This can be a challenging task for large nanostructures. Computational techniques and software have been developed to tackle this problem. Cadnano [1] is a widely employed software created to assist with the design of lattice-based DNA nanostructures. It is highly reliable, as it constrains the cross-section of the design to two lattice types, square and honeycomb, to ensure the proper placement of the crossovers and therefore high folding percentages of the DNA nanostructure *in vitro*. However, it is not straightforward to create nanostructures mixing different types of lattices, it does not provide means to a modular approach and all DNA double helices must be parallel in a design (Figure 1a), which reduces the design possibilities. For example, alternative DNA nanostructure concepts such as wireframe DNA origamis are difficult to realize in Cadnano. In addition, automated design workflows using geometric structures as input and an appropriate visualization of the designs is missing. This represents a significant challenge with the increasing complexity of nanostructures.

**Figure 1:**
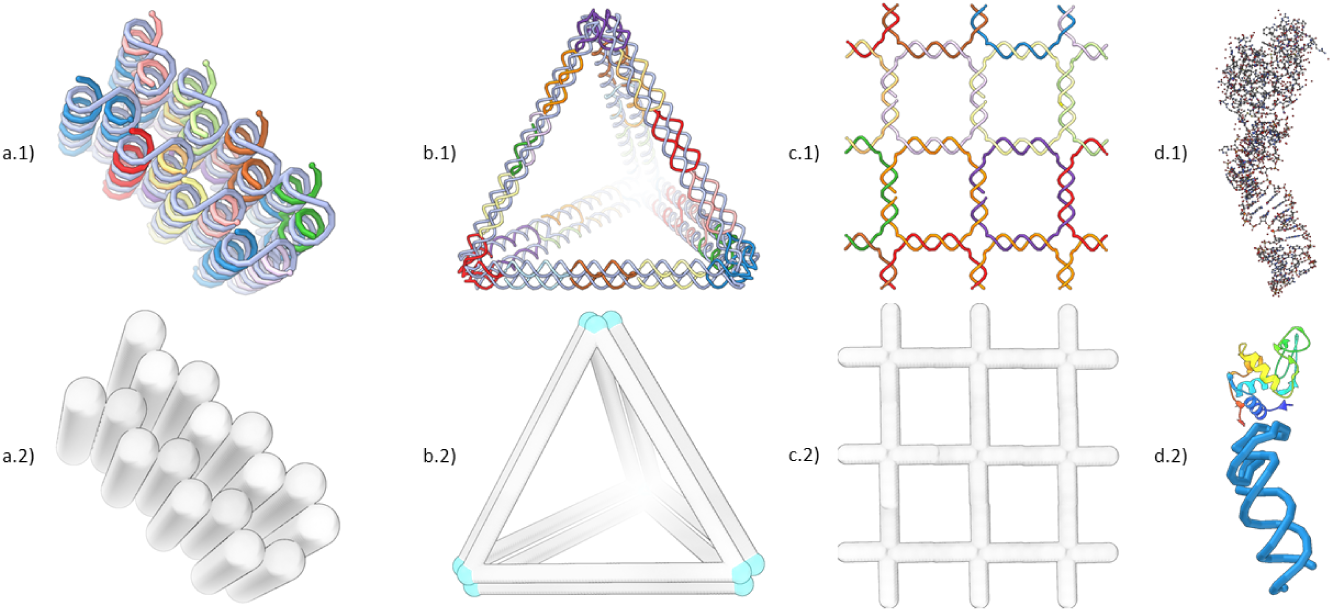
Screenshots of different DNA nanostructure design concepts and their visualization in Adenita. (a) A multi-layer DNA brick designed with Cadnano, (b) a tetrahedron designed using Daedalus and (c) a squared lattice DNA Tile that was manually designed. In Cadnano structures, all double helices must be parallel, while DNA wireframe approaches place double strands at the edges of a mesh. DNA Tiles require the repetition of one or several small DNA motifs through space to create a 2D or 3D shape, in this case we used a four-arm Holliday junction. (d.1) shows the PDB 4M4O, formed by a protein and an aptamer, loaded with the default all-atom models, in (d.2) a combination of Adenita’s visual model and Samson’s secondary structure visualization makes possible to simplify the representation of such molecule.

In contrast, automated design workflows for wireframed DNA nanostructures were developed. To design wireframed DNA origami objects, the target shape is represented as a graph and the problem of tracing the scaffold through is, then, the known NP-complete problem of finding an A-trail along a graph. VHelix [4] is a pipeline of tools that provide a semi-manual interface. Its input is a triangular mesh whose edges are partially represented by either one or two double helices (to allow for the routing of the scaffold). It outputs the sequence of the staples for the target wireframe, as well as a model that can be loaded into the 3D-modeling software Maya [12], allowing an inspection after the nanostructure model has been created. Daedalus [5] provides a completely automated tool that can work with non-triangular meshes too, in which edges are always represented by two double strands (Figure 1b). Nevertheless, it does not provide interactive methods to make *a posteriori* changes and the final structure has to be inspected using all-atom models with external tools.

Another approach to the design of DNA nanostructures are DNA tiles. It is a modular strategy that employs small motifs with sticky ends that can be used to create higher order 2D and 3D nanostructures (Figure 1c). DNA tiles have been particularly used for the self-assembly of periodic structures, such as 2D lattices [13] and 3D crystals [14], and recently it has also been employed for constructing random complex shapes [15]. To the best of our knowledge, no specific software has been created so far to design nanostructures using arbitrary DNA tiles.

With the aforementioned applications, the field of DNA nanotechnology advances rapidly and the involved DNA nanostructures are ever increasing in size and complexity [16]. With the recent developments in hybrid DNA-protein systems [17, 18], the need for sophisticated modeling and visualization tools becomes apparent. We aim to facilitate the combination of DNA nanostructures with other molecules, such as aptamers, proteins or nanoparticles, in a feasible manner that does not require a large pipeline of tools or the inspection of nanostructures at the atomic scale.

In this work we present Adenita, an interactive 3D tool for the design, visualization and modification of DNA nanostructures, independently of the chosen paradigm. We provide a semi-manual and highly modular approach that is well-suited, not only for multilayer or wireframe DNA origami approaches, but also for the use of DNA tiles. We have developed a hierarchical data model that is able to describe arbitrary DNA nanostructures and a sophisticated multiscale visualization method that depicts the nanostructures on multiple levels of detail allowing the user to operate on a desired level of detail for a specific task. Furthermore, real-time feedback of structural stability is integrated into the visual model.

Through simple 3D interactions and visibility handling, different components or parametrizable predefined structures can be interactively loaded, created or combined into higher-order structures. A straightforward application of this approach allows the user to import Cadnano designs, make free-form designs of DNA tiles or create wireframe nanostructures using the Daedalus algorithm, and therefore combine different approaches *in silico*. Furthermore, we have developed Adenita as a plugin for SAMSON [19], a software for adaptive 3D modeling and simulation of nanosystems, making it possible to edit and work on arbitrary DNA nanostructures while also visualizing and editing other systems, such as aptamers or proteins (Figure 1d).

Adenita has been developed to overcome the design limitations of the existing DNA origami design tools, with a focus on the modeling of nanostructures in more realistic molecular environments, which will significantly facilitate the prediction of intended and unintended interactions. Our contribution can be summarized as:

- Integration across folding patterns: A unified DNA nanostructure framework that integrates all major folding strategies and allows their smooth combination
- Integration along scales of conceptual organization: A unified modeling concept that seamlessly integrates a wide spectrum of semantic scales on which one can study and manipulate the nanostructure
- Multi-stage DNA-nanotechnology self-assembly: A convincing use case in which elementary pieces can be created in one stage and in consecutive stages they can be integrated together to form a more complex design.

## 2 MATERIALS AND METHODS

### 2.1 Dependencies and hardware requirements

We implemented Adenita as a suite of plugins for the computational nanoscience software SAMSON [19]. Adenita enables the user to create, modify and visualize DNA-based structures.

We allow for an optional integration with *ntthal* from the Primer3 package [20] to compute thermodynamic parameters of the nanostructure. Adenita has been developed with the help of Boost [21] and Rapidjson [22] libraries.

To generate wireframe nanostructures we employ the Daedalus algorithm [5].

A graphics card is highly recommended in order to guarantee interactive framerates and a smooth visualization of the 3D structures.

### 2.2 Experimental methods

The DNA nanostructures were prepared based on protocols already described in the work of Ahmadi et al. [23, 24]. These nanorods were initially designed with Cadnano 2.2.0 using the p8064 single stranded scaffold. Subsequently, they were modified with Adenita to form a new nanostructure in a cross shape composed of two nanorods. Each protruding and invading strand necessary to form the crosses was assigned to one of the two nanorods composing the design. Individual nanorods were self-assembled separately with a 1:10 scaffold to staple strand ratio in the Tris buffer (TB) solution (5*mM* Tris, 1*mM* EDTA, 5*mM NaCl*) containing 18*mM MgCl*_2_. Annealing was performed by exposing the reaction mixture to 65*°C* for 15 min and then cooling down from 65 to 25*°C* by 1*°C* every 40 min in a one-day thermal ramp. The nanorods were purified using the PEG precipitation method based on a protocol described by Evi Stahl et al [25]. In brief, 100*µl* of the nanostructure sample (in TB including 18*mM MgCl*_2_) was mixed with an equivalent volume of 22*mM MgCl*_2_ supplemented TB (100*µl*), followed by the addition of 200*µl* of purification buffer (15% (w/v) PEG 8000, 5*mM* Tris, 1*mM* EDTA and 505*mM NaCl*). The solution was then mixed gently by the tube inversion and spun at 16 000g at r.t. for 25 min. The supernatant was then carefully discarded, and the pellet was dissolved in the TB buffer supplemented with 16 mM *MgCl*_2_, followed by incubation for one day at r.t. at 650 rpm. For super-assembly of crosses, the stoichiometric amount of purified nanorods were mixed, followed by incubation overnight at r.t. at 700 rpm.

TEM images of the crosses and individual rods were obtained by diluting samples with origami buffer 1:10 and vortexing for 10*s*. Diluted samples were negatively stained using uranyl acetate on 300-mesh carbon coated grids that had been glow discharged for 40*s* and imaged on an FEI Tecnai T12 Spirit electron microscope. Images were collected at a nominal magnification of 1650x using a defocus of 25 to 40*µm*. Fiji [26, 27] was used to analyze the TEM images.

## 3 RESULTS

### 3.1 Description of the software

Adenita describes arbitrary DNA nanostructures using a data model that comprises two related parent-child hierarchies.

The first hierarchy describes single-stranded DNA. The top element is the single-strand, whose children are the nucleotides, ordered from the 5’ to the 3’ end. Nucleotides are formed by a backbone and a sidechain, which in turn group the atoms.

The second hierarchy describes the geometry of the DNA nanostructure. It is based on a graph model where the double strands are the edges that compose the target geometry. This model is straightforward in the case of wireframe nanostructures [4, 5], but it can also be applied to any rasterized target shape. We can then consider the edges or double strands as the top element whose children are the base pairs that form them. The base pairs can be generalized to also include unpaired regions and motifs, such as the poly-T regions of Daedalus designs illustrated at the vertices in blue in the Figure 1b.

The relationship between both hierarchies is established through the nucleotides composing each base pair (Figure 2f). It is determined by the routing of the scaffold and the placement of the staples, which can be done manually by the user or with the help of algorithms, such as Daedalus.

**Figure 2:**
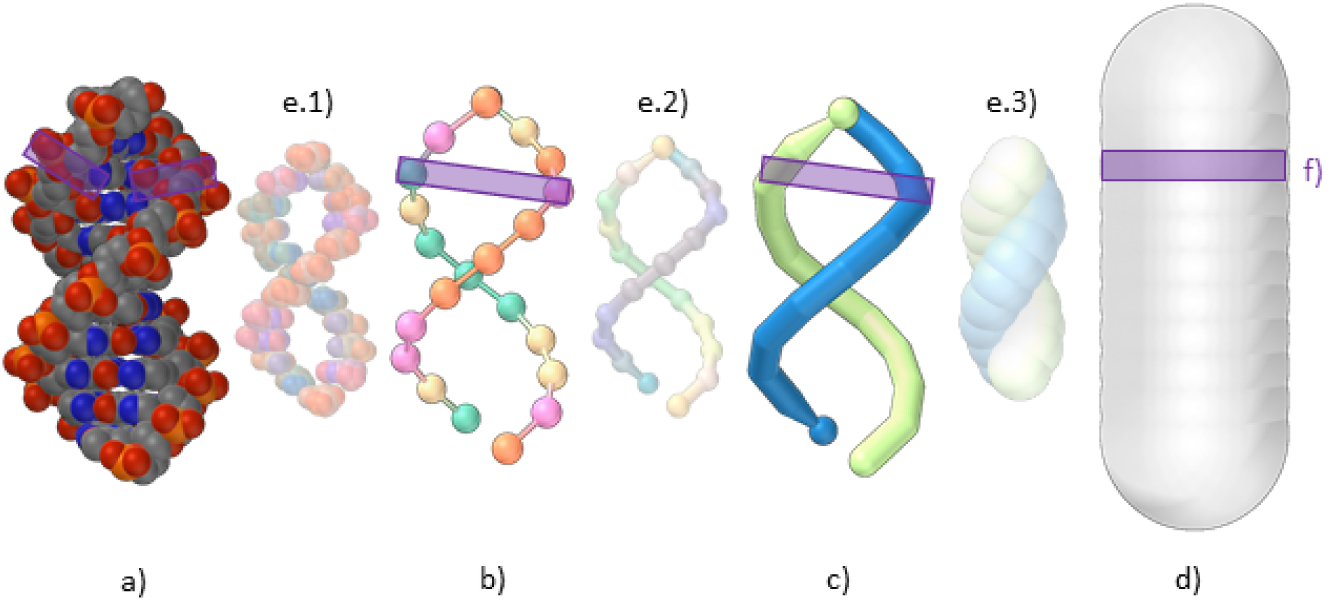
The data model describes every nucleotide using its backbone and side-chain positions fetched from its all-atom representation (a). A single strand is represented as a chain of nucleotides with directionality (b). Double strands can be represented as paired regions of single strands (c) or as the segments that trace the target shape (d). The visual model represents graphically all scales of the data model and allows for a seamlessly transition between them (e). The bottom-up scales (a and b) are related to the top scale (d) through the positioning of the base pairs (f).

Adenita estimates the position of nucleotides using a top-down approach. Once the geometry of the target shape has been specified, the positioning of base pairs and therefore nucleotides is inferred using a model based on B-DNA and idealized base pairs [28].

Our visualization concept, depicts the DNA nanostructure in various forms of details, which we call scales. In our multiscale approach [29], the user can continuously transition between multiple scales and related atomic representation to high-level double stranded representation (Figure 2). This multiscale provides users with means to operate in any desired scale and expect the results to be represented in other scales. For better compatibility with 2D designs and Cadnano, the visualization includes a multidimensional approach [30], which provides a 2D and 1D view for Cadnano designs.

Modeling of DNA nanostructures results in an idealized representation of the object that can be experimentally realized. We have implemented a highlight mode which provides immediate feedback to the design process, helping to visually detect interesting patters in the design, such as single-strands with specific lengths, unassigned bases or crossovers. It is also possible to employ *ntthal*, from the Primer3 suite [20] to calculate thermodynamic parameters of the binding regions on demand. A binding region is defined as consecutive base pairs that are not limited by a strand end or a crossover. All analysis results are color coded in the visualization. It is also possible to control the visibility of all elements of the data model. By controlling the scale, highlight mode and visibility the user can tailor the visualization to be better suited for a specific task.

The combination of the data structure, DNA model and visualization makes it possible to create, visualize, modify and analyze DNA nanostructures. The output of these processes can be a re-usable model of the DNA nanostructure, the list of sequences needed for the *in vitro* self-assembly or structural files for simulations in oxDNA [31]. Basic modifications include deletion of various elements, concatenation or insertion of DNA strands and breaking strands by deleting the phosphodiester bond between nucleotides, amongst others.

Users can access all modifications, editors and options through an intuitive user interface. Through parametrized editors, the user can choose predefined shapes, and then vary some parameters to create a customized version of the selected shape. Some shapes provide basic building blocks, like the drawing of simple DNA strands or non-routed nanotubes. Others can provide more complex shapes, such as the wireframed editor, which allows the user to select a target 3D geometry and modify some of its parameters in a controllable manner while visualizing it, after which Daedalus is used to produce the DNA nanostructure.

### 3.2 DNA nanostructure manipulation

The editors allow the modification of existing DNA nanostructures, as well as the creation of new ones from scratch. It is possible to add single and double strands, straight or circular, cut any strand by either deleting the phosphodiester bond between nucleotides or deleting a nucleotide, and reconnect different strands (Figure 3). To connect strands with each other, it is possible to move them in close proximity or to introduce a new strand to link them.

**Figure 3:**
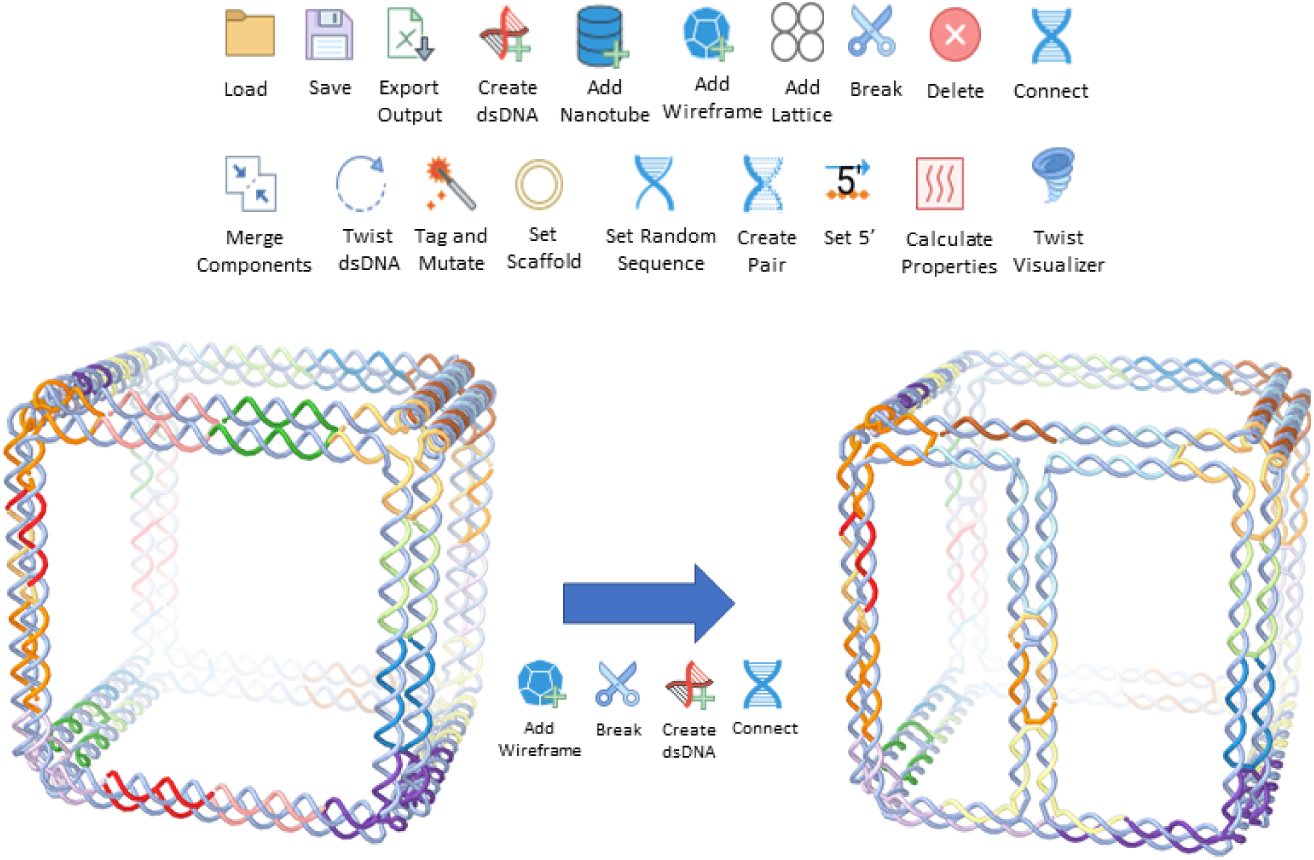
Depiction of the different editors for the interactive modeling of DNA nanostructures. Here we demonstrate how a wireframe cube created with the Daedalus algorithm can be further edited to include an extra edge. In this way, it is possible to introduce *in silico* internal faces to Daedalus designs by creating extra edges on the proper faces, overcoming one of its limitations.

To showcase these editors and evaluate the precision of our data model, we designed cross-shaped nanostructures comprising two individual multilayer DNA origami nanorods. The nanorods consists of around 16,000 nucleotides, have an approximate size of 350*nm* × 8*nm* × 4*nm* (Figure 4a) and were originally designed for other applications [23].

**Figure 4:**
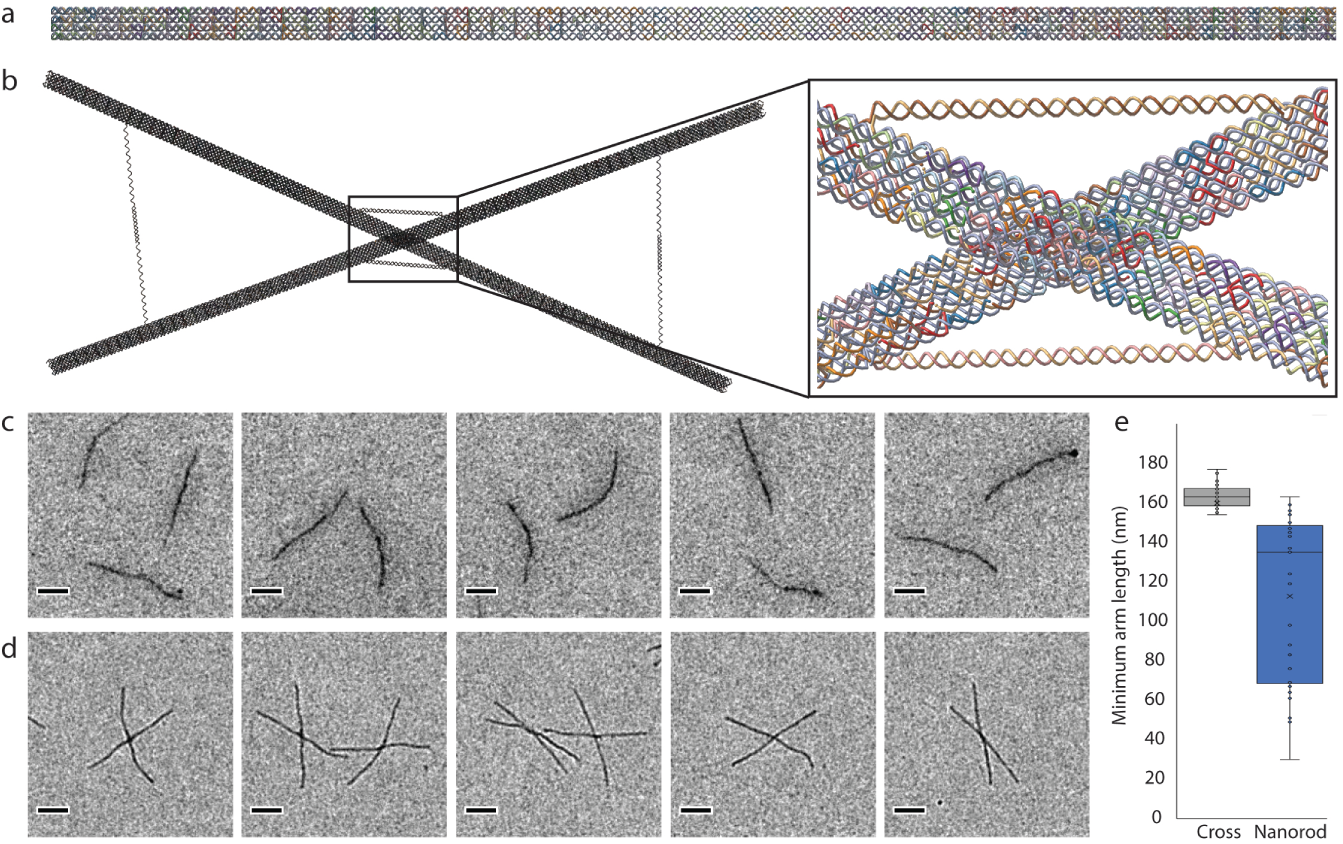
The DNA origami nanorod *(a)* and the cross *(b)* as designed in Adenita. *(c)* Negative stain TEM micrographs taken of the control nanorods. *(d)* Negative stain TEM images of the cross. *(e)* Some crosses appear to be formed when nanorods superimpose each other on the slide, a statistical analysis of the cross arms length was performed to check that crosses were correctly folded.

We used the nanorod to create a simple cross. Each cross consists of two nanorods that were imported into Adenita as separate components, and connected at different points with invading and extruding strands with the help of editors (Figure 4a and 4b). Additional strands to constrain the cross angles and to give further stability were also added. We estimated with the help of the visual representation the connection points and the lengths of the new strands.

Both single nanorods and crosses were imaged with TEM using negative stain preparation (Figure 4c and 4d). Due to superimposition of nanorods on the control slide that resulted in structurally deviating crosses, we measured the length of the arms of all observed crosses. In the case of the designed crosses, we expected certain regularity on the arms length, as by design each arm should be around half the nanorod length, i.e. 175*nm*. In the case of crosses appearing on the TEM images of the controls, we expected to see greater variation on the arms length. These experiments confirmed that our data model allows a realistic in silico manipulation of long DNA nanostructures (Figure 4e).

## 4 DISCUSSION

Computational support to model 3D objects has become standard in many areas due to facilitating the design and fabrication process. The publication of Cadnano boosted the emerging DNA nanotechnology field providing researchers a simple tool to create nanoscale multilayer DNA origami objects. Due to limitations associated with the multilayer design concept (e.g. structures cannot fold at physiological ion concentrations) and the software tool itself, alternative design tools such as Deadalus and vHelix were developed promoting new design concepts. However, these tools were also limited to a single DNA origami design concept and, in contrast to Cadnano, were lacking an appropriate user-interface and intuitive manual modeling possibilities. Adenita was developed to overcome these limitations. In addition, it addresses the increasing complexity of DNA nanostructures and their envisioned applications allowing the incorporation of structural data from pdb-files into the modelling. Beside the possibility to create complex objects such as DNA nanostructure-based artificial enzymes [18], we used Adenita for the design of biosensor surfaces, DNA pores and a DNA robot (supplementary information).

The long nanorods were selected to showcase another powerful feature of Adenita. In general, the precise modeling of DNA nanostructures becomes more challenging as the nanostructures grow in size due to the lack of accurate structural prediction. Thus, an imprecise model introduces an error at every helix turn, which increases as the helix becomes longer. In some cases, this can be overcome by using nanostructures that have been extensively evaluated in the laboratory or with simulations. We took advantage of such a modular approach when designing the DNA origami crosses, as we had previously tested the nanorods. An alternative approach to overcome this problem are simulations that estimate a more realistic *in silico* model. For this purpose, we have implemented an export function for oxDNA simulations of the nanostructure model. However, in the future our DNA model could be fine tuned using experimental data. More detailed spatial information on the e.g. helix turns will further improve the nanostructure designs. Nevertheless, the experiment with the crosses demonstrated that the implemented model is precise enough to modify large structures.

Future work will also involve optimizing performance, so Adenita is capable of working more smoothly with larger designs or with the new Gigadalton structures. This could be achieved by incorporating a representation of the nanostructure at the Gigadalton scale, or modifying the visualization to handle global and local representations.

Adenita is not only a framework capable of handling different paradigms, but also introduces novel concepts to the design of DNA nanostructures, such as a modular approach, a novel visualization and an environment capable of handling also other types of molecules, such as proteins or aptamers. We have shown that Adenita is capable of handling large structures and that the combination of its data model and the novel visualization gives the user the ability to edit and visualize nanostructures effectively. It combines in one tool several steps of current DNA nanostructure design pipelines. The use of several scales in the data model as well as in the visualization allows the user to work with the DNA nanostructure at different resolutions in parallel. Furthermore, this can be combined with editors and analysis options, extending the design possibilities much further than any other existing tool. At the same time we recognize the strengths of the current methods, and we have found a way to incorporate them into Adenita’s work-flow.

We foresee that the combination of an user-friendly environment with a modular approach will prompt a sharing-economy approach in the DNA nanotechnology community.

## 5 SOFTWARE AVAILABILITY

Adenita is open-source and publicly available. It can be downloaded through SAMSON’s Elements store for free (https://www.samson-connect.net/elements.html). Source code can be found at:

- https://github.com/edellano/Adenita-SAMSON-Edition-Win- (Windows)
- https://github.com/edellano/Adenita-SAMSON-Edition-Linux (Linux)

## Supporting information

User Documentation

Showcase designs files

## 6 AUTHOR CONTRIBUTIONS

E. de Llano: Was responsible for the software concept and design. Developed the data model, most of the backend and part of the frontend. Designed the crosses on Adenita. Did the data analysis on the TEM images. Wrote most of the paper. Created figures. Reviewed the paper.

H. Miao: Was responsible for the visualization and interaction concept. Developed the visualization, part of the backend and most of the frontend. Created user documentation. Created figures. Wrote and reviewed the paper.

Y. Ahmadi: Was responsible for the design of nanorods in Cadnano and the in vitro folding of the nanostructures. Wrote and reviewed the paper. Created figures.

A. J. Wilson and M. Beeby: Were responsible for the TEM imaging. Wrote and reviewed the paper.

I. Viola: Worked on the visualization concept. Reviewed the paper.

I. Barisic: Tested the software. Idealized the crosses. Ideliazed and designed the biosensor surface, DNA pore and DNA robot on Adenita. Wrote and reviewed the paper. Created figures.

## 7 ACKNOWLEDGMENTS

This work has received funding from the European Union’s Horizon 2020 research and innovation program under grant agreement No 686647, from WWTF under the ILLVISATION grant (VRG11-010) and from the King Abdullah University of Science and Technology (KAUST) Office of Sponsored Research (OSR) under Award No. OSR-2019-CPF-410.

We want to thank all Adenita beta users for their feedback.

## References

[1] Douglas, S. M. and Marblestone, A. H. and Teerapittayanon, S. and Vazquez, A. and Church, G. M. and Shih, W. M. (2009) Rapid prototyping of 3D DNA-origami shapes with caDNAno. Nucleic Acids Res., 37, 5001–5006.

[2] Rothemund, P. (2006) Folding DNA to create nanoscale shapes and patterns. Nature, 440, 297–302.

[3] Douglas, S. M. and Dietz, H. and Liedl, T. and Högberg, B. and Graf, F. and Shih, W. M. (2009) Self-assembly of DNA into nanoscale three-dimensional shapes Nature, 10.1038/nature08016

[4] Benson, E. and Mohammed, A. and Gardell, J. and Masich, S. and Czeizler, E. and Orponen, P. and Högberg, B. (2015) DNA rendering of polyhedral meshes at the nanoscale. Nature, 523, 441–444.

[5] Veneziano, R. and Ratanalert, S. and Zhang, K. and Zhang, F. and Yan, H. and Chiu, W. and Bathe, M. (2016) Designer nanoscale DNA assemblies programmed from the top down. Science, 10.1126/science.aaf4388.

[6] Nickels, P. C. and Wünsch, B. and Holzmeister, P. and Bae, W. and Kneer, L. M. and Grohmann, D. and Tinnefeld, P. and Liedl, T. (2016) Molecular force spectroscopy with a DNA origami-based nanoscopic force clamp. Science, 10.1126/science.aah5974

[7] Linko, V. and Eerikäinen, M. and Kostiainen, M. A. (2015) A modular DNA origami-based enzyme cascade nanoreactor Chemical Communications, 10.1039/C4CC08472A

[8] Bell, N. A.W. and Engst, C. R. and Ablay, M. and Divitini, G. and Ducati, C. and Liedl, T. and Keyser, U. F. (2012) DNA Origami Nanopores Nano Lett., 12, 512–517.

[9] Selnihhin, D. and Sparvath, S. M. and Preus, S. and Birkedal, V. and Andersen, E. S. (2018) Multi-Fluorophore DNA Origami Beacon as a Biosensing Platform ACS Nano, 10.1021/acsnano.8b01510

[10] Andersen, E. S. and Dong, M. and Nielsen M. M. and Jahn, K. and Subramani, R. and Mamdouh, W. and Golas M. M. and Sander, B. and Stark, H. and Oliveira, C. L. P. and Pedersen, J. S. and Birkedal, V. and Besenbacher, F. and Gothelf, K. V. and Kjems, J. (2009) Self-assembly of a nanoscale DNA box with a controllable lid Nature, 459, 73–76.

[11] Ke, Y. and Sharma, J. and Liu, M. and Jahn, K. and Liu, Y. and Yan, H. (2009) Scaffolded DNA Origami of a DNA Tetrahedron Molecular Container Nano Lett., 9, 2445–2447.

[12] Maya: 3D computer animation, modeling, simulation, and rendering software. (2019). Retrieved from: https://www.autodesk.com/products/maya/overview

[13] Winfree, E. and Liu, F. and Wenzler, L. A. and Seeman, N. C. (1998) Design and self-assembly of two-dimensional DNA crystals. Nature, 10.1038/28998

[14] Zheng, J. and Birktoft, J. J. and Chen, Y. and Wang, T. and Sha, R. and Constantinou, P. E. and Ginell, S. L. and Mao, C. and Seeman, N. C. (2009) From molecular to macroscopic via the rational design of a self-assembled 3D DNA crystal. Nature, 10.1038/nature08274

[15] Wei, B. and Dai, M. and Yin, P. (2012) Complex shapes self-assembled from single-stranded DNA tiles. Nature, 10.1038/nature11075

[16] Wagenbauer, K. F. and Sigl, C. and Dietz, H. (2017) Gigadalton-scale shape-programmable DNA assemblies Nature, 10.1038/nature24651

[17] Kosuri, P. and Altheimer, B. D. and Dai, M. and Yin, P. and Zhuang, X. (2019) Rotation tracking of genome-processing enzymes using DNA origami rotors Nature, 10.1038/s41586-019-1397-7

[18] Kekic, T. and Ahmadi, Y. and Barisic, I. (2019) Enzyme Catalytic Activity Emulated Within DNA-based Nanodevice bioRxiv, 10.1101/804518

[19] SAMSON, the open molecular modeling platform. (2019). Retrieved from: https://www.samson-connect.net

[20] Untergasser, A. and Cutcutache, I. and Koressaar, T. and Ye, J. and Faircloth, B. C. and Remm, M. and Rozen, S. G. (2012) Primer3–new capabilities and interfaces. Nucleic Acids Res., 10.1093/nar/gks596

[21] Boost [Computer library]. (2019). Retrieved from: https://www.boost.org/

[22] Rapidjson [Computer library]. (2019). Retrieved from: http://rapidjson.org/

[23] Ahmadi, Y. and De Llano, E. and Barisic, I. (2018) (Poly)Cation-Induced Protection of Conventional and Wireframe DNA Origami Nanostructures. Nanoscale, 10.1039/C7NR09461B

[24] Ahmadi, Y. and Barisic, I. (2019) Gene-therapy Inspired Polycation Coating for Protection of DNA Origami Nanostructures. J. Vis. Exp., 10.3791/58771

[25] Stahl, E. and Martin, T. G. and Praetorius, F. and Dietz, H. (2014) Facile and scalable preparation of pure and dense DNA origami solutions. Angew. Chem. Int. Ed. Engl., 10.1002/anie.201405991

[26] Schindelin, J. and Arganda-Carreras, I. and Frise, E. and Kaynig, V. and Longair, M. and Pietzsch, T. and Preibisch, S. and Rueden, C. and Saalfeld, S. and Schmid, B. and Tinevez, J. and White, D. J. and Hartenstein, V. and Eliceiri, K. and Tomancak P. and Cardona, A. (2012) Fiji: an open-source platform for biological-image analysis Nature, 10.1038/nmeth.2019

[27] Rueden, C. T. and Schindelin, J. and Hiner, M. C. and DeZonia, B. E. and Walter, A. E. and Arena, E. T. and Eliceiri, K. W (2017) ImageJ2: ImageJ for the next generation of scientific image data BMC Bioinform., 10.1186/s12859-017-1934-z

[28] Lu, X. and Olso, W. K. (2003) 3DNA: a software package for the analysis, rebuilding and visualization of three-dimensional nucleic acid structures Nucleic Acids Res., 10.1093/nar/gkg680

[29] Miao, H. and De Llano, E. and Sorger, J. and Ahmadi, Y. and Kekic, T. and Isenberg, T. and Gröller, M. E. and Barisic, I. and Viola, I. (2018) Multiscale Visualization and Scale-Adaptive Modification of DNA Nanostructures. IEEE Trans. Vis. Comput. Graph., 10.1109/TVCG.2017.2743981

[30] Miao, H. and De Llano, E. and Isenberg, T. and Gröller, M. E. and Barisic, I. and Viola, I. (2018) DimSUM: Dimension and Scale Unifying Maps for Visual Abstraction of DNA Origami Structures. Comput. Graph. Forum, 10.1111/cgf.13429

[31] Sulc, P. and Romano, F. and Ouldridge, T. E. and Rovigatti, L. and Doye, J. P. K. and Louis, A. A. (2012) Sequence-dependent thermodynamics of a coarse-grained DNA model. J. Chem. Phys., 10.1063/1.4754132

