## Supplementary Information for "Adenita: Interactive 3D modeling and visualization of DNA Nanostructures"

Supplementary material

### SEQUENCES

#### Scaffold

The scaffold employed to fold the structures was a single-stranded scaffold DNA type p8064 with sequence:

GGCAATGACCTGATAGCCTTTGTAGATCTCTCAAAAATAGCTACCCTCTCCGGCATTAATTTATCAGCTAGAACGGTTGAATATCATATTGATGGTGATTTG
ACTGTCTCCGGCCTTTCTCACCCTTTTGAATCTTTACCTACACATTACTCAGGCATTGCATTTAAAATATATGAGGGTTCTAAAAATTTTTATCCTTGCGTTGA
AATAAAGGCTTCTCCCGCAAAAGTATTACAGGGTCATAATGTTTTTGGTACAACCGATTTAGCTTTATGCTCTGAGGCTTTATTGCTTAATTTTGCTAATTCTT
TGCCTTGCCTGTATGATTTATTGGATGTTAATGCTACTACTATTAGTAGAATTGATGCCACCTTTTCAGCTCGCGCCCCAAATGAAAATATAGCTAAACAGGT
TATTGACCATTTGCGAAATGTATCTAATGGTCAAACTAAATCTACTCGTTCGCAGAATTGGGAATCAACTGTTATATGGAATGAAACTTCCAGACACCGTAC
TTTAGTTGCATATTTAAAACATGTTGAGCTACAGCATTATATTCAGCAATTAAGCTCTAAGCCATCCGCAAAAATGACCTCTTATCAAAAGGAGCAATTAAA
GGTACTCTCTAATCCTGACCTGTTGGAGTTTGCTTCCGGTCTGGTTCGCTTTGAAGCTCGAATTAAAACGCGATATTTGAAGTCTTTCGGGCTTCCTCTTAAT
CTTTTTGATGCAATCCGCTTTGCTTCTGACTATAATAGTCAGGGTAAAGACCTGATTTTTGATTTATGGTCATTCTCGTTTTCTGAACTGTTTAAAGCATTTGA
GGGGGATTCAATGAATATTTATGACGATTCCGCAGTATTGGACGCTATCCAGTCTAAACATTTTACTATTACCCCCTCTGGCAAAACTTCTTTTGCAAAAGCC
TCTCGCTATTTTGGTTTTTATCGTCGTCTGGTAAACGAGGGTTATGATAGTGTTGCTCTTACTATGCCTCGTAATTCCTTTTGGCGTTATGTATCTGCATTAGT
TGAATGTGGTATTCCTAAATCTCAACTGATGAATCTTTCTACCTGTAATAATGTTGTTCCGTTAGTTCGTTTTATTAACGTAGATTTTTCTTCCCAACGTCCTGA
CTGGTATAATGAGCCAGTTCTTAAAATCGCATAAGGTAATTCACAATGATTAAAGTTGAAATTAAACCATCTCAAGCCCAATTTACTACTCGTTCTGGTGTTT
CTCGTCAGGGCAAGCCTTATTCACTGAATGAGCAGCTTTGTTACGTTGATTTGGGTAATGAATATCCGGTTCTTGTCAAGATTACTCTTGATGAAGGTCAGC
CAGCCTATGCGCCTGGTCTGTACACCGTTCATCTGTCCTCTTTCAAAGTTGGTCAGTTCGGTTCCCTTATGATTGACCGTCTGCGCCTCGTTCCGGCTAAGTA
ACATGGAGCAGGTCGCGGATTTCGACACAATTTATCAGGCGATGATACAAATCTCCGTTGTACTTTGTTTCGCGCTTGGTATAATCGCTGGGGGTCAAAGA
TGAGTGTTTTAGTGTATTCTTTTGCCTCTTTCGTTTTAGGTTGGTGCCTTCGTAGTGGCATTACGTATTTTACCCGTTTAATGGAAACTTCCTCATGAAAAAGT
CTTTAGTCCTCAAAGCCTCTGTAGCCGTTGCTACCCTCGTTCCGATGCTGTCTTTCGCTGCTGAGGGTGACGATCCCGCAAAAGCGGCCTTTAACTCCCTGCA
AGCCTCAGCGACCGAATATATCGGTTATGCGTGGGCGATGGTTGTTGTCATTGTCGGCGCAACTATCGGTATCAAGCTGTTTAAGAAATTCACCTCGAAAG
CAAGCTGATAAACCGATACAATTAAAGGCTCCTTTTGGAGCCTTTTTTTTGGAGATTTTCAACGTGAAAAAATTATTATTCGCAATTCCTTTAGTTGTTCCTTT
CTATTCTCACTCCGCTGAAACTGTTGAAAGTTGTTTAGCAAAATCCCATACAGAAAATTCATTTACTAACGTCTGGAAAGACGACAAAACTTTAGATCGTTAC
GCTAACTATGAGGGCTGTCTGTGGAATGCTACAGGCGTTGTAGTTTGTACTGGTGACGAAACTCAGTGTTACGGTACATGGGTTCCTATTGGGCTTGCTAT
CCCTGAAAATGAGGGTGGTGGCTCTGAGGGTGGCGGTTCTGAGGGTGGCGGTTCTGAGGGTGGCGGTACTAAACCTCCTGAGTACGGTGATACACCTAT
TCCGGGCTATACTTATATCAACCCTCTCGACGGCACTTATCCGCCTGGTACTGAGCAAAACCCCGCTAATCCTAATCCTTCTCTTGAGGAGTCTCAGCCTCTT
AATACTTTCATGTTTCAGAATAATAGGTTCCGAAATAGGCAGGGGGCATTAACTGTTTATACGGGCACTGTTACTCAAGGCACTGACCCCGTTAAAACTTAT
TACCAGTACACTCCTGTATCATCAAAAGCCATGTATGACGCTTACTGGAACGGTAAATTCAGAGACTGCGCTTTCCATTCTGGCTTTAATGAGGATTTATTTG
TTTGTGAATATCAAGGCCAATCGTCTGACCTGCCTCAACCTCCTGTCAATGCTGGCGGCGGCTCTGGTGGTGGTTCTGGTGGCGGCTCTGAGGGTGGTGGC
TCTGAGGGTGGCGGTTCTGAGGGTGGCGGCTCTGAGGGAGGCGGTTCCGGTGGTGGCTCTGGTTCCGGTGATTTTGATTATGAAAAGATGGCAAACGCT
AATAAGGGGGCTATGACCGAAAATGCCGATGAAAACGCGCTACAGTCTGACGCTAAAGGCAAACTTGATTCTGTCGCTACTGATTACGGTGCTGCTATCGA
TGGTTTCATTGGTGACGTTTCCGGCCTTGCTAATGGTAATGGTGCTACTGGTGATTTTGCTGGCTCTAATTCCCAAATGGCTCAAGTCGGTGACGGTGATAA
TTCACCTTTAATGAATAATTTCCGTCAATATTTACCTTCCCTCCCTCAATCGGTTGAATGTCGCCCTTTTGTCTTTGGCGCTGGTAAACCATATGAATTTTCTAT
TGATTGTGACAAAATAAACTTATTCCGTGGTGTCTTTGCGTTTCTTTTATATGTTGCCACCTTTATGTATGTATTTTCTACGTTTGCTAACATACTGCGTAATAA
GGAGTCTTAATCATGCCAGTTCTTTTGGGTATTCCGTTATTATTGCGTTTCCTCGGTTTCCTTCTGGTAACTTTGTTCGGCTATCTGCTTACTTTTCTTAAAAAG
GGCTTCGGTAAGATAGCTATTGCTATTTCATTGTTTCTTGCTCTTATTATTGGGCTTAACTCAATTCTTGTGGGTTATCTCTCTGATATTAGCGCTCAATTACCC
TCTGACTTTGTTCAGGGTGTTCAGTTAATTCTCCCGTCTAATGCGCTTCCCTGTTTTTATGTTATTCTCTCTGTAAAGGCTGCTATTTTCATTTTTGACGTTAAA
CAAAAAATCGTTTCTTATTTGGATTGGGATAAATAATATGGCTGTTTATTTTGTAACTGGCAAATTAGGCTCTGGAAAGACGCTCGTTAGCGTTGGTAAGAT
TCAGGATAAAATTGTAGCTGGGTGCAAAATAGCAACTAATCTTGATTTAAGGCTTCAAAACCTCCCGCAAGTCGGGAGGTTCGCTAAAACGCCTCGCGTTC
TTAGAATACCGGATAAGCCTTCTATATCTGATTTGCTTGCTATTGGGCGCGGTAATGATTCCTACGATGAAAATAAAAACGGCTTGCTTGTTCTCGATGAGT
GCGGTACTTGGTTTAATACCCGTTCTTGGAATGATAAGGAAAGACAGCCGATTATTGATTGGTTTCTACATGCTCGTAAATTAGGATGGGATATTATTTTTC
TTGTTCAGGACTTATCTATTGTTGATAAACAGGCGCGTTCTGCATTAGCTGAACATGTTGTTTATTGTCGTCGTCTGGACAGAATTACTTTACCTTTTGTCGG
TACTTTATATTCTCTTATTACTGGCTCGAAAATGCCTCTGCCTAAATTACATGTTGGCGTTGTTAAATATGGCGATTCTCAATTAAGCCCTACTGTTGAGCGTT
GGCTTTATACTGGTAAGAATTTGTATAACGCATATGATACTAAACAGGCTTTTTCTAGTAATTATGATTCCGGTGTTTATTCTTATTTAACGCCTTATTTATCA
CACGGTCGGTATTTCAAACCATTAAATTTAGGTCAGAAGATGAAATTAACTAAAATATATTTGAAAAAGTTTTCTCGCGTTCTTTGTCTTGCGATTGGATTTG
CATCAGCATTTACATATAGTTATATAACCCAACCTAAGCCGGAGGTTAAAAAGGTAGTCTCTCAGACCTATGATTTTGATAAATTCACTATTGACTCTTCTCA
GCGTCTTAATCTAAGCTATCGCTATGTTTTCAAGGATTCTAAGGGAAAATTAATTAATAGCGACGATTTACAGAAGCAAGGTTATTCACTCACATATATTGA
TTTATGTACTGTTTCCATTAAAAAAGGTAATTCAAATGAAATTGTTAAATGTAATTAATTTTGTTTTCTTGATGTTTGTTTCATCATCTTCTTTTGCTCAGGTAA
TTGAAATGAATAATTCGCCTCTGCGCGATTTTGTAACTTGGTATTCAAAGCAATCAGGCGAATCCGTTATTGTTTCTCCCGATGTAAAAGGTACTGTTACTGT
ATATTCATCTGACGTTAAACCTGAAAATCTACGCAATTTCTTTATTTCTGTTTTACGTGCAAATAATTTTGATATGGTAGGTTCTAACCCTTCCATTATTCAGA
AGTATAATCCAAACAATCAGGATTATATTGATGAATTGCCATCATCTGATAATCAGGAATATGATGATAATTCCGCTCCTTCTGGTGGTTTCTTTGTTCCGCA
AAATGATAATGTTACTCAAACTTTTAAAATTAATAACGTTCGGGCAAAGGATTTAATACGAGTTGTCGAATTGTTTGTAAAGTCTAATACTTCTAAATCCTCA

AATGTATTATCTATTGACGGCTCTAATCTATTAGTTGTTAGTGCTCCTAAAGATATTTTAGATAACCTTCCTCAATTCCTTTCAACTGTTGATTTGCCAACTGAC
CAGATATTGATTGAGGGTTTGATATTTGAGGTTCAGCAAGGTGATGCTTTAGATTTTTCATTTGCTGCTGGCTCTCAGCGTGGCACTGTTGCAGGCGGTGTT
AATACTGACCGCCTCACCTCTGTTTTATCTTCTGCTGGTGGTTCGTTCGGTATTTTTAATGGCGATGTTTTAGGGCTATCAGTTCGCGCATTAAAGACTAATA
GCCATTCAAAAATATTGTCTGTGCCACGTATTCTTACGCTTTCAGGTCAGAAGGGTTCTATCTCTGTTGGCCAGAATGTCCCTTTTATTACTGGTCGTGTGAC
TGGTGAATCTGCCAATGTAAATAATCCATTTCAGACGATTGAGCGTCAAAATGTAGGTATTTCCATGAGCGTTTTTCCTGTTGCAATGGCTGGCGGTAATAT
TGTTCTGGATATTACCAGCAAGGCCGATAGTTTGAGTTCTTCTACTCAGGCAAGTGATGTTATTACTAATCAAAGAAGTATTGCTACAACGGTTAATTTGCG
TGATGGACAGACTCTTTTACTCGGTGGCCTCACTGATTATAAAAACACTTCTCAGGATTCTGGCGTACCGTTCCTGTCTAAAATCCCTTTAATCGGCCTCCTG
TTTAGCTCCCGCTCTGATTCTAACGAGGAAAGCACGTTATACGTGCTCGTCAAAGCAACCATAGTACGCGCCCTGTAGCGGCGCATTAAGCGCGGCGGGTG
TGGTGGTTACGCGCAGCGTGACCGCTACACTTGCCAGCGCCCTAGCGCCCGCTCCTTTCGCTTTCTTCCCTTCCTTTCTCGCCACGTTCGCCGGCTTTCCCCGT
CAAGCTCTAAATCGGGGGCTCCCTTTAGGGTTCCGATTTAGTGCTTTACGGCACCTCGACCCCAAAAAACTTGATTTGGGTGATGGTTCACGTAGTGGGCC
ATCGCCCTGATAGACGGTTTTTCGCCCTTTGACGTTGGAGTCCACGTTCTTTAATAGTGGACTCTTGTTCCAAACTGGAACAACACTCAACCCTATCTCGGGC
TATTCTTTTGATTTATAAGGGATTTTGCCGATTTCGGAACCACCATCAAACAGGATTTTCGCCTGCTGGGGCAAACCAGCGTGGACCGCTTGCTGCAACTCT
CTCAGGGCCAGGCGGTGAAGGGCAATCAGCTGTTGCCCGTCTCACTGGTGAAAAGAAAAACCACCCTGGCGCCCAATACGCAAACCGCCTCTCCCCGCGC
GTTGGCCGATTCATTAATGCAGCTGGCACGACAGGTTTCCCGACTGGAAAGCGGGCAGTGAGCGCAACGCAATTAATGTGAGTTAGCTCACTCATTAGGC
ACCCCAGGCTTTACACTTTATGCTTCCGGCTCGTATGTTGTGTGGAATTGTGAGCGGATAACAATTTCACACAGGAAACAGCTATGACCATGATTACGAATT
CGAGCTCGGTACCCGGGGATCCTCAACTGTGAGGAGGCTCACGGACGCGAAGAACAGGCACGCGTGCTGGCAGAAACCCCCGGTATGACCGTGAAAACG
GCCCGCCGCATTCTGGCCGCAGCACCACAGAGTGCACAGGCGCGCAGTGACACTGCGCTGGATCGTCTGATGCAGGGGGCACCGGCACCGCTGGCTGCA
GGTAACCCGGCATCTGATGCCGTTAACGATTTGCTGAACACACCAGTGTAAGGGATGTTTATGACGAGCAAAGAAACCTTTACCCATTACCAGCCGCAGGG
CAACAGTGACCCGGCTCATACCGCAACCGCGCCCGGCGGATTGAGTGCGAAAGCGCCTGCAATGACCCCGCTGATGCTGGACACCTCCAGCCGTAAGCTG
GTTGCGTGGGATGGCACCACCGACGGTGCTGCCGTTGGCATTCTTGCGGTTGCTGCTGACCAGACCAGCACCACGCTGACGTTCTACAAGTCCGGCACGTT
CCGTTATGAGGATGTGCTCTGGCCGGAGGCTGCCAGCGACGAGACGAAAAAACGGACCGCGTTTGCCGGAACGGCAATCAGCATCGTTTAACTTTACCCT
TCATCACTAAAGGCCGCCTGTGCGGCTTTTTTTACGGGATTTTTTTATGTCGATGTACACAACCGCCCAACTGCTGGCGGCAAATGAGCAGAAATTTAAGTT
TGATCCGCTGTTTCTGCGTCTCTTTTTCCGTGAGAGCTATCCCTTCACCACGGAGAAAGTCTATCTCTCACAAATTCCGGGACTGGTAAACATGGCGCTGTAC
GTTTCGCCGATTGTTTCCGGTGAGGTTATCCGTTCCCGTGGCGGCTCCACCTCTGAAAGCTTGGCACTGGCCGTCGTTTTACAACGTCGTGACTGGGAAAAC
CCTGGCGTTACCCAACTTAATCGCCTTGCAGCACATCCCCCTTTCGCCAGCTGGCGTAATAGCGAAGAGGCCCGCACCGATCGCCCTTCCCAACAGTTGCGC
AGCCTGAATGGCGAATGGCGCTTTGCCTGGTTTCCGGCACCAGAAGCGGTGCCGGAAAGCTGGCTGGAGTGCGATCTTCCTGAGGCCGATACTGTCGTCG
TCCCCTCAAACTGGCAGATGCACGGTTACGATGCGCCCATCTACACCAACGTGACCTATCCCATTACGGTCAATCCGCCGTTTGTTCCCACGGAGAATCCGA
CGGGTTGTTACTCGCTCACATTTAATGTTGATGAAAGCTGGCTACAGGAAGGCCAGACGCGAATTATTTTTGATGGCGTTCCTATTGGTTAAAAAATGAGCT
GATTTAACAAAAATTTAATGCGAATTTTAACAAAATATTAACGTTTACAATTTAAATATTTGCTTATACAATCTTCCTGTTTTTGGGGCTTTTCTGATTATCAAC
CGGGGTACATATGATTGACATGCTAGTTTTACGATTACCGTTCATCGATTCTCTTGTTTGCTCCAGACTCTCA

### List of staples

| Common sequences to the nanorod and the cross | |
|---|---|
| 1 | ATCAAAAAGATTAAGATTCGCGAGCTTTC |
| 2 | GTGGACTCCAACGTCGGAGTGTACCGGAACG |
| 3 | AGGCTATCAGGTCATTTCCATATTATACC |
| 4 | GCGGTCACGCTGCGCGTGGAAAGCCCCAATAA |
| 5 | TTTATTTTTCAAGAAACAAAAGAAGATGATGANNNNN |
| 6 | AACGCCATCAAAAATAAGGAAGCCGCTTTTGC |
| 7 | AGAGCACATCCTCATAGTACAACGCCCCCTGC |
| 8 | TTAATATTTTGTTAAAAGCAAACTGTAAGAGC |
| 9 | CAGCAGGCGAAAATCCAGACTCCTATACACTA |
| 10 | AAAGGAATCATAATTAAACCCACAAGAGAAAGAGGACAGTTAGAAG |
| 11 | NNNNNNNCGTAGGAATCACATTTGGGAATNNNNNNN |
| 12 | ATTCAACCGTTCTAGCATTTCGCAAGATGGTT |
| 13 | GTCGCTATTAATTAATTATCCCATATCAAGAT |
| 14 | CATTCCACAGACAGCCGAGCCGGACGGTGCCC |
| 15 | GTTTGAAATACCGACTATGCGTGAGGGTA |
| 16 | NNNNNNNTTTAAACATGACCGTAATGGNN |
| 17 | TTTTTCTTTTCACCAGATAGGTGTACTAAAGA |
| 18 | GATTCCCAATTCTGCGTTTGAGAGAACAATCG |
| 19 | GAGGCGGTTTGCGTATTACCGCCAGAGGGTAG |
| 20 | ACACCAACTATTAGACTTTACAAAAGCCGTCAGAGTTAAGGCAGTCTC |
| 21 | TTGAAAACATAGCGATACAATAGAACAATTTT |
| 22 | CCGGCTTAGGTTGGGTAGAATATATCCAAATA |
| 23 | CTATTTCGGAACCTATCCCTTATATGCCGGAC |
| 24 | CAGGCGGATAAGTGCCGCCTGGCCAACCAGCT |
| 25 | GCAGAAGAATATAAATTCATAATCAAAATC |
| 26 | TAGCGTTTTTAACCGTCCGAACGAACCACCA |

| | |
|---|---|
| **27** | TATAATCAGTGAGGCCCAGAGCCAAAATACAT |
| **28** | TGAGCCTCCTCACAGTTGTTTCAGCTTGCGAAT |
| **29** | NNNNNNNGGAACAACACGCGTGCCTGTNN |
| **30** | ATAACCTGTTTAGCTATCACCATCCTCCGTGG |
| **31** | TTCAACGCAAGGATAATTAGCATCATCAA |
| **32** | CAGGAGGCCGATTAACACCACCCGAATACCC |
| **33** | ATTGTATAAGCAAATAACCTTTAATACATAAC |
| **34** | ACCGCCACCCTCAGAAGGCCAACGTCACTGTT |
| **35** | GGCGCGGTTGCGGTATATCGGAACCCCTCAGA |
| **36** | TAGTAAATGAATTTTCAATCATGGTGCGGCCA |
| **37** | TGGTAATATCCAGAAGTAATCAGCAACCGAT |
| **38** | TCCTGTAGCCAGCTTTAAGCAAAGTAATAGTA |
| **39** | CTTTCCTCGTTAGAATACCAGAGCAGAAGGAA |
| **40** | ATTGGCCTTGATATTCGCTACAGGAAGGTTAT |
| **41** | TACTTCTTTGATTAGTCATTTTCGTTTGTCAC |
| **42** | TTTCAACAGAGGATCCGTCATACCGGGGGTTT |
| **43** | TCAGGATTAGAGAGTTTTAAATCTCTTCG |
| **44** | TTCTTACCAGTATAAATGACCTAATAAAAGTT |
| **45** | TGCCAGTTGCGGGCCGTGAAAATATTTTGCTAAACAAC |
| **46** | TGCCAGTTCAGACGACGACAATAAACAACATGAGTGAATTCATATCAA |
| **47** | TGGCACAGCGACATTTAGCGACAGAATCAA |
| **48** | NNNNNNNCATTGGCAGATTCACCTGAAATGGATTATTTANNNNNNN |
| **49** | ACCGTCGGTGGTGCCAACTACGAACTCAGTAC |
| **50** | GGGTTACCTGCAGCCCGATAGTCCTGTAG |
| **51** | GCATTGACAGGAGGTAGCACGTAGTTGGCAA |
| **52** | ACTCAGGAGGTTTAGTGGGCGCCATCCGCCG |
| **53** | GAGCGAGTAACAACCATCAGGTAATACTG |
| **54** | NNNNNNNTAGAGCCAGCAAAATCACCCAATCGTCAGTCACAC |
| **55** | TATGGTTGCTTTGACGTGAGGCAGAAAAGTAA |
| **56** | ACGAGCATGTAGAAACCTTGCTTCACATCGGG |
| **57** | TGACTATTATAGTCAGCATCAACAAGTATCGG |
| **58** | NNNNNNAACAAACACATNNNNN |
| **59** | TGATAATCAGAAAAGTTTTGCGATTTAGG |
| **60** | CTGCATTAATGAATCCCGCCACCTCACCCTC |
| **61** | TGTCCATCACGCAAAGCCATCTTAGAAACGC |
| **62** | ATTGCGTAATCGCCATAAGTTTATGTCATAGCCCCCTTAT |
| **63** | CAACATGTTTTAAATATGAACGACGACGG |
| **64** | AGTGCCCGTATAAACGGTTGAGTCTCCGGCC |
| **65** | AGTTAATTTCATCTTCGCCAACGCGGAGAATT |
| **66** | CGAGCTCGAATTCGTTGTATGGGCTCCAAAA |
| **67** | GGATTAGGATTAGCGCACGCTGGACGGCAGC |
| **68** | TTCAAATATCGCGTTTTTTTTTAACCAGGCAA |
| **69** | AGCCTGTTTAGTATCACGTGTGATATTAAATC |
| **70** | CGCCACCAGAACCACCCAGAGCGGAAATATCA |
| **71** | AAACCAAGTACCGCATTCATTTAGAGGCG |
| **72** | TCGTCACCAGTACAAATGCCTAATCAGATGCC |
| **73** | CCGGAGACAGTCAAATATTTTCAAACACC |
| **74** | GTTGCAGCAAGCGGTCGGGTTTTGGGCACCAA |
| **75** | TCATATATTTTAAATGACATCCAAATTCATTA |
| **76** | TAACTCACATTAATTGACCGTAACCATAACCG |
| **77** | ATTCCACACAACATACCTCATAGTTGAATTTC |

| | |
|---|---|
| **78** | GAATAGCCCGAGATAGAGTTAATGGAGATTTG |
| **79** | GCAAACAAGAGAATCGATGCAACTCTACGTTA |
| **80** | CTGACCAACTTTATTATAGATAACAAGTGTA |
| **81** | TTTTTTAATGGAAACATCCAAGAATATTCTAA |
| **82** | GTGTCTGGAAGTTTCATGCCTGAGAGCCGCCA |
| **83** | TCGAGCCAGTAATAAGTATATAACTTCATCAA |
| **84** | ACGCCAACATGTAATTCCAATCGCCATCATAT |
| **85** | GAACAAGAAAAATAATTTCCCTGTCAGAT |
| **86** | GACCAGTAATTTCAATCCCAATAGGAACAAGCAAGCCGTT |
| **87** | TAAGTTTTAACGGGGTTCCACTATCGGTCCGT |
| **88** | TAAGAGCACTAATATCCAACTCGTAAATAAGGCGTTAAATAAGAATAA |
| **89** | GCAGCACCCAATATTAAAGCGTAAGAATACG |
| **90** | GGCTTTTGATGATACAAAAGGGCGTAAACGAT |
| **91** | CCGCCACCCTCAGAGCAGGGATTTAAAAATCT |
| **92** | AATGCTGATGCAAATTAGGCAGAAAAATG |
| **93** | TCATAGGTCTGAGAGAATTCTGTCACAAAATA |
| **94** | CATTGCAACAGGAAAAACGTCACCCGGAAATT |
| **95** | ACCGGAACACCGAGTAAGGTGAGGCGGTCAGTAAGGGTTA |
| **96** | AATTACATTTAACAATCTCATCGACAAGCAAA |
| **97** | GAATACCAAACCCTTCAATATTGAAATGAAACCATCGATA |
| **98** | CCACCACCCTCATTTTAAACCTGTAGGTTTCT |
| **99** | TTTGATAAGAGGTCATCCCCAAAAGGATGTGC |
| **100** | CTGCCAGCTAAAGGAAGGAGTGAGAATAGAAANNNNNN |
| **101** | ACAAAAGGTAAAGTACTACCTTACTTCTG |
| **102** | TCCTGATTCACCTTGCAATAACGTCAGAGC |
| **103** | AAAGCATATCAGATTTAACGTCAGGCATTT |
| **104** | AAACACTCTAGTAAATTAGACTTTAATATGAT |
| **105** | AACAGCCATGTTAGCACGCCTGCGTGTTTT |
| **106** | TAATTTCACGAAAGAGAATGCCATTTGCCC |
| **107** | TTGCTCGTCCAGCTGGAATGCAGATTGCTCCT |
| **108** | AACACTATACAACCATAACGGCATGAGTGAGC |
| **109** | CCAGTGCCGCGCTTTCCTTTGAGGATCACCGT |
| **110** | TTTCCAGAGTGGCAACTAAAACAGAAAGAGTC |
| **111** | GCTGATTTAAAAAAGCTGACCTAAATTAAG |
| **112** | ATATATTGAATTACGTGCGGGCTGTAAACG |
| **113** | AACTGAACCGAAGCCCAATTGAGGGCGCGTAC |
| **114** | GCAGATAGAAGCGCTTTGCGGATATATTTT |
| **115** | AAAAGAACTTTTTTGTGATGGCAATATATGTA |
| **116** | TATGCGATGTAATGCCTCCCACGCCTGAGAGA |
| **117** | TACGGCTGATAACCCTGGCTCATAACAGTT |
| **118** | GCCAAAGAGGTTTTGATACCTTTTTGTAAATC |
| **119** | GTTTGCCTAAACTATCTTTGAATG |
| **120** | TAAAATACTTTAAGAATCACCGGAATCTACAA |
| **121** | AATAATGGATTAACACAACGTAGACCACCGGA |
| **122** | AAAAGGCTAGACTGGAACGACGACTTAAATGT |
| **123** | TGCAAGGCGGCTGGTAGGGATCGCTCAGAG |
| **124** | GAACGCGAGGAAGGTATGACCTGACCGCCAGC |
| **125** | ATTCATTAGAAGGCAATCGCGCGAATTACC |
| **126** | GTGAATAAGAAACAAAACGGAACGAATCAAAA |
| **127** | TAAAGGCCATTCAACTCGAAAGGGACAGGAAG |
| **128** | CCTGCATCCTTCTGGTAGCGAGAGCGAAAGAC |

| | |
|---|---|
| **129** | AAATGTTTCCAAAAGGTGTGGTGCTCATAGCT |
| **130** | AGAGATAGAGTTACAATTATCCGGCGGGTATT |
| **131** | AACCCTCAGAAGGAGCCAGAGAGAATTTAACA |
| **132** | CCAAGCGCGGCTTGCCCACGGAAAGAGAAAGG |
| **133** | CGGCACCGAGACGATCTTTCGAGGTAGCGTAA |
| **134** | CGGAATTTACCGCAAGGCAAAAGACAAGAGAA |
| **135** | ACATAAAGGCCTAATTGAACCTACTATCAAAA |
| **136** | AATACCACGCTTTTGCATGGGTAACGTGCCAG |
| **137** | TATAATCCGAGCCAGCTTAAGACTCCTCAGAA |
| **138** | AGGCGCAGATAGGCTGATCCCGTAGCCTTTAT |
| **139** | ACTTGAGCTTACCGCGTACCTGAGACAAAATT |
| **140** | GAGTAATCCTCCATGTGGCAAACGTAAAGAAC |
| **141** | GAATGCGTGAGGGGTAGCGTCCCTTTACCC |
| **142** | TTGCGGGACAAAAGGGACAATATTGGCCTTGC |
| **143** | AGAAACGATGGCATGAAGCAAATGTAGACAGG |
| **144** | TTGTAGATAGCTCTCTGACGAGATTTGGGG |
| **145** | AAAATAGCAACGCAATTGAACCTCGAGCTAAA |
| **146** | CCTAAAACGATTTTCAACCCAGCTTAAGTCCT |
| **147** | CTTTTTCACGTTGGGAAGAGGTGGAGTCTGGA |
| **148** | TGAGTAACTTGAAAGGTTTTTAAGGTCAGACG |
| **149** | TTAAACAGACCAAAATGCCGGAAACCAATAGG |
| **150** | TCATTTGCCTCGTCGCAATTGTGTTGAGTAAC |
| **151** | ATCCTGAACACGGAATTAAAAATATGTAGCAA |
| **152** | TTGGGAAGAAATCGTTCGCCCACGACTGAGTT |
| **153** | ACCGAGGAAGCCTTTAGGAATTATAAGACAAA |
| **154** | AGCAGCGAAGATTCATTTGGGTAAACCCCGGT |
| **155** | CTAAAATGAACGTTCAAAGTCATATACAAA |
| **156** | TATCTTACACCCTGAAATTAATTTATTTAATG |
| **157** | ATCAACAGATTATCATATTAGACGTCAACAGT |
| **158** | CAGCAGCAGTGAGAGATGGGCTTGAATGGTCA |
| **159** | CATTGCAGAAGCTTTCAGAAAAATAAAGTACG |
| **160** | CAACGGCAACTAACGTTGTAAAGTAATCGT |
| **161** | AAAAGAAGCAGCTTGCCAGCGCAGCTCACA |
| **162** | TGTTCAGCGGCGATCGGAGGCATACCAACAGG |
| **163** | GCTATTAGAGTAACAGAGCCTTAACCTAATTT |
| **164** | ATAAAACGTACAGAGGGCACTCAAGGGTGG |
| **165** | CCTCAGGAGTGCACTCAGCCTTTACCAGACGT |
| **166** | TAGATTAGCAATTCGAAGAGAGATCTAGAAAA |
| **167** | GCCCTGCGATTAAGCAGTTGAGGATGGCTT |
| **168** | CGGAATCGTTTCACGTTTTTCACGCCGGGTAC |
| **169** | CCAGGCGCACGGTCAATCATAAGGTCATACAT |
| **170** | AAAACAGGGCCGAACAAAGTTACCCGCCGCCA |
| **171** | CTCATTAAAGCCAGAATAACCACCACAACTAA |
| **172** | AGCGAACCAGACCGGAATTCGCATCGCAACTG |
| **173** | AAGTATTAAGAGGCTGTGTTTGATGGTCTGGT |
| **174** | ACGCGCCTGTTTATCAAGCTTAGAATAAAGAA |
| **175** | TTGACCATTAGATACTGATAAACCAGTCC |
| **176** | TTAGCAAGGCCGGAAACGCTCATTGGCCAAC |
| **177** | GGTTTATTTTTGCCAGAGGGGGCGGATTGC |
| **178** | CGCAGTATATTATTTGGATTATTTTAACCT |
| **179** | GCGCGTTTTCATCGGAATAACATCTGATAGC |

| | |
|---|---|
| **180** | AATAGGAACCCATGTCGTTGCGCCACTGGTG |
| **181** | TGACCATAAATCAAAACGTCGGATCGTGCATC |
| **182** | ATAGTAGTAGCATTACAATGCCTTTCTGC |
| **183** | AGGTAGAAAAGACAGCGAGCCGGGCGCGGGGA |
| **184** | ATAAGTATAGCCCGGATGAGACGGAGCGGGGT |
| **185** | TGTCTTTCCTTATCATGTACATAAATTGCTTT |
| **186** | CTTTGCCCATCTTTAGGAGCACTAACACCCGC |
| **187** | AGGGCTTAATTGAGATTTCAAAACAAAGA |
| **188** | AATTATTCATAAAAGGGACATTCGGAAATA |
| **189** | GAATATACTCTTTAATGCGCGAACACTTGC |
| **190** | CCTAAAAACTTTAACATGTTTATTAATGCC |
| **191** | AAAGACACTCTTACCAAAACAGAATTAAGACG |
| **192** | TATCATCAACGTAAAACTTAAATGAGTAAT |
| **193** | AGAGCTTAATTGCTGATCATATGTCGCCAGGG |
| **194** | CGATCTAAAGTTTTGTGTTATCCGTGTCACT |
| **195** | TGAGGGAGGGCGTTTTTTCGCCTGATCAATAT |
| **196** | ACCGCCTCCCTCAGACCTGAGAAAACAGTGC |
| **197** | AATAATTTTCATAAATATCGTAACTCTCCGTG |
| **198** | CAATGACACATAACCCTCAGGCTGTAAATTTT |
| **199** | CAATAAAGCCTCAGAGCGGGAGAAAAAAAAGC |
| **200** | CTATTACGCATAAACATCCCTTATCACTGC |
| **201** | TGAAGGGAACGTCAGCGTGGTGCTGGTGGTTC |
| **202** | CGGGAACGGGAGGTGTCCAGCATCGCAACAGC |
| **203** | GGAGAGGGTAGCTATTAACGAGTAAATTACCT |
| **204** | CCTACATTTTGACGCTAGTAGCACCGTCACCG |
| **205** | ATGTGAGTGAATAACCAATCAATCCCGAC |
| **206** | AACCACCAATCAATATCTGGTCATAACGTG |
| **207** | CTGAGTAGAAGAACTCTTAGCGTCTTACCAGC |
| **208** | GTGTAGGTAAAGATTCGGCATCAACTCATTCA |
| **209** | GGAACAAACGGCGGATGTTCAGAAAATCCCCCTCAAATGCNNNN |
| **210** | CGCACAGGCGGCCGATATGAACGGTAAATCGGTTGTACC |
| **211** | CGAAATCGGCAAAATTATTCTGACGATTATA |
| **212** | CGCGCTTAATGCGCCACAAACAAGCAATAGC |
| **213** | TTTTCCCAGTCACGACGGAACAACCTGTAGCT |
| **214** | CACGCTGATGATTGTTTATCCCAAAAGTACCG |
| **215** | TGATTGCCCTTCACCGTCGAGAGTAAACGGG |
| **216** | CCGCTTTCCAGTCGGGCAGGGATACAGGGAGT |
| **217** | TGTTAAATCAGCTCATAATTCGCAGACGA |
| **218** | CTGAGAAGAGTCAATTTCAGCTGAGCGTC |
| **219** | GCGCGCCTAGATCGCACTCCAGCCTCTGGCCT |
| **220** | ATTATGACCCTGTAATACTTTTGCATAAAGCTGTACAGA |
| **221** | AGGTCACGTTGGTGTAGATGGGCGCATTCATTGAACGAGAA |
| **222** | GCGAAACGTACAGCGCTCATTGTGGATTTAGT |
| **223** | AACGGTACGCCAGAATGCCGCCACCCTTATTA |
| **224** | AGAACGAGATCTTTGACCCCCAGAACATGA |
| **225** | ATTGAGCGAGAAACAATGAAATAATAAATC |
| **226** | AAAACTAGCATGTCAAATATAATGATTATTAC |
| **227** | AGCGCCATTCGCCATTCGTTTACAGCTTCAA |
| **228** | GTTTCCTGTGTGAAATTCGTCTTTATTGTATC |
| **229** | CAGAAACAGCGGATCACAAAGCTGTTCTACTA |
| **230** | AGTCAGGATGAGGAAGTTTCCATGGTTGAT |

| | |
|---|---|
| **231** | AATTATTTGCACGTAACGCTAACAATGCAGA |
| **232** | GAACGCGAGAAAACTTATCGCCATATAACATA |
| **233** | AGAAACAATAACGGAAGCGAACCTAATCGGC |
| **234** | GCCAAAAGCGGTCGCTGAGGCTTGGCAAGCCC |
| **235** | TCAGATATAAAGGTGAATTATCACCATTACCA |
| **236** | TAGTTGCTAGAAAATTCATATGGTAGACTGTA |

| **Nanorod sequences** | |
|---|---|
| **1** | CGCGAGCTGAAAAGGTAAAAGGGTAAGAGACG |
| **2** | ACAGGCAAGGCAAAGAAAAATTTTGTTGTGTA |
| **3** | TGAATTTACCGTTCCGCGCTGGTACATTTGAGTTTAGTGATGAAG |
| **4** | GTGTAAAGCCTGGGGCTACAACGTGCGCCGA |
| **5** | GGTAAAGTAAAAACCGTCTATCAGGCGCTAGGAGTAAGCGGAACCGAA |
| **6** | CAGTTTGGAACAAGAGCAGTGCCTCGAAATCC |
| **7** | CATCGACAGCCGTTCCTACTTAGCTGGTAA |
| **8** | AATCAATATTTTGCGGTTTAACTAGAATCC |
| **9** | CCCAAATCGCCTGATATGGCAGCGTTGTTC |
| **10** | TTTTTCGTCGCCAGCAAACCGGATTAAATCAT |
| **11** | GCGACCTGTTGACAAGGTTGGGCGTAGAACCC |
| **12** | CGATAAAACTTGATACAGCGGTGCAGCATAAA |

| **Cross sequences** | |
|---|---|
| **1** | TCATTTGCCTCGTCGCAATTGTAAGTTTACAAGAGCAGTGCCTCGAAATCC |
| **2** | CGATAAAACTTGATACAGCGGTGCAGCATAAAGTG |
| **3** | CGAAATCGGCAAAATTTCATCAGAATGGGTTACTATTGCGTCTGGAACAATAATACATTCAATCGACG GCATACCCAAAATATTCGTCATGTAATATCCTTGCCTTGCATGGGACTATGGCGTAAGCGCGCTGGTA CATTTGAGTTTAGTGATGAAG |
| **4** | TTTTTCGTCGCCAGCAAACCGGATTAAATCATTTTGAAAAGCAAACCGGATTAAATCAT |
| **5** | CATCGACAGCCGTTCCTACTTAGCTGGTAATTTCAAACATCGACAGCCGTTCCTACTTAGCTGGTAA |
| **6** | ACACCACATCTCCAATGTGCACTTTGCTAACTTGAGGACTTAGATATTAATAGATGTGCTATTAGCTTT CGTTTATCTGTTAACTGCGACGTTGAATGATTGGGATGCCATAATGAAGGGTAAGAGACG |
| **7** | ACCACAGCATATCCTCCGTTCCTAGTGGCGTTTCTGCATTCTATCGCAGAATAACATTAAGATAGTCTA AGAATGCAGTTGTCCCCCCGTTAAAGTATATAATGCACGATGTCCTAGGCCAGCCGTTAAGTGATGTT AGAATCTAGTTCCGTCCCCACTTTAAAGCCTGGGGCTACAACGTGCGCCGA |
| **8** | AATCAATATTTTGCGGTTTAACTA |
| **9** | CAGTTTGGAACAAGAGCAGTGCCTCGAAATCCTTTGAAAGGCGTAGAACCC |
| **10** | TGAATTTACCGTTCCAGTAAGCGGAACCGAA |
| **11** | CCAACCTGTGACACACCGAACCGTGGATCTGTATTTATTCAAAGTAATATAACGCAATGTTTAAATTTT GCTTTACACCTTTTTTAAGGTATGTAAGACTCGCATGCAGTTATGGCCGAGAAGGTTCAACTCTGAATA GTCTATATAACTTCGCGCGAGGGAATCC |

DESIGN SHOWCASES

### DNA nanopores

Nanopore composed of six double strands

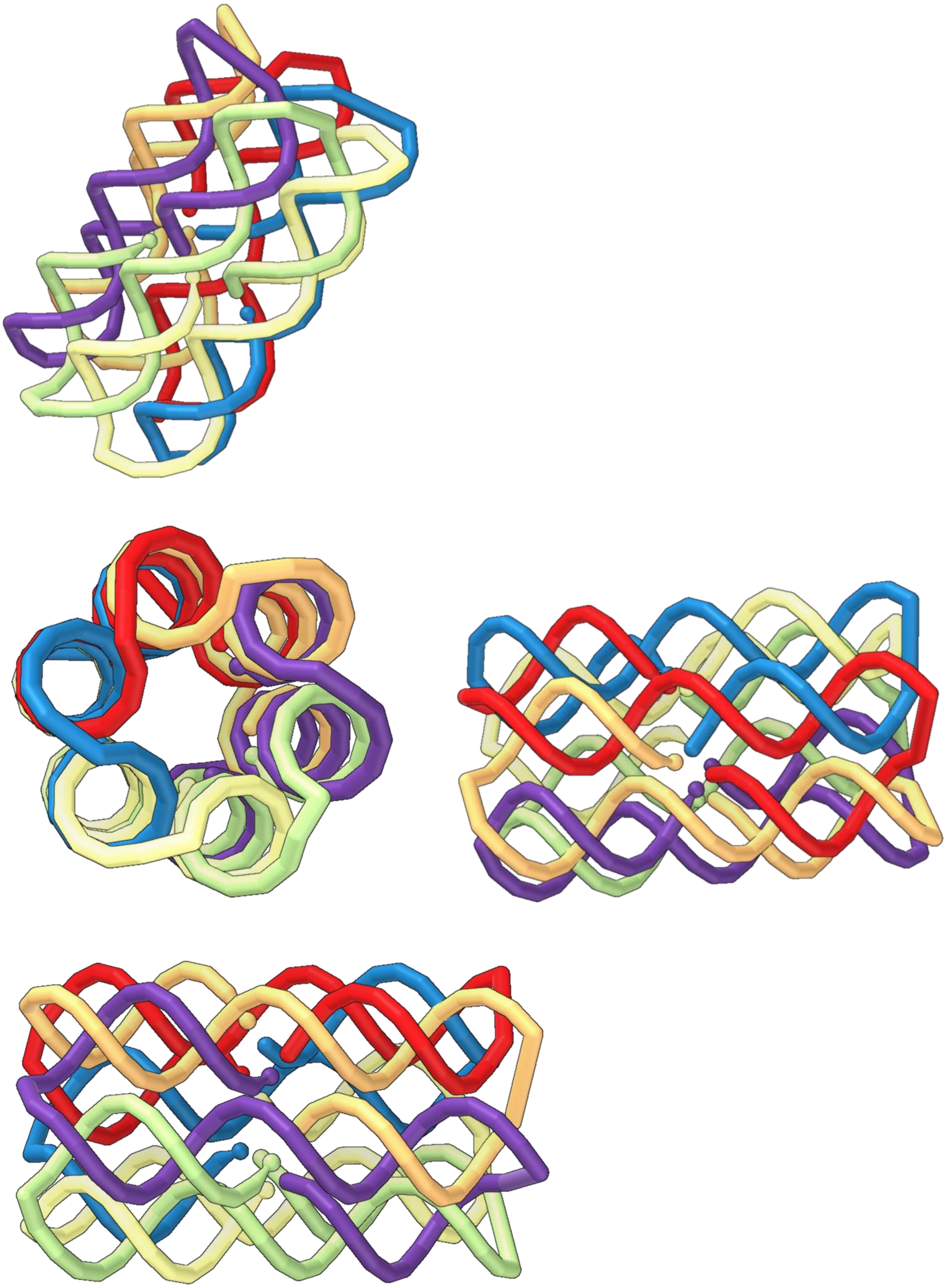

Nanopore composed of twelve double strands

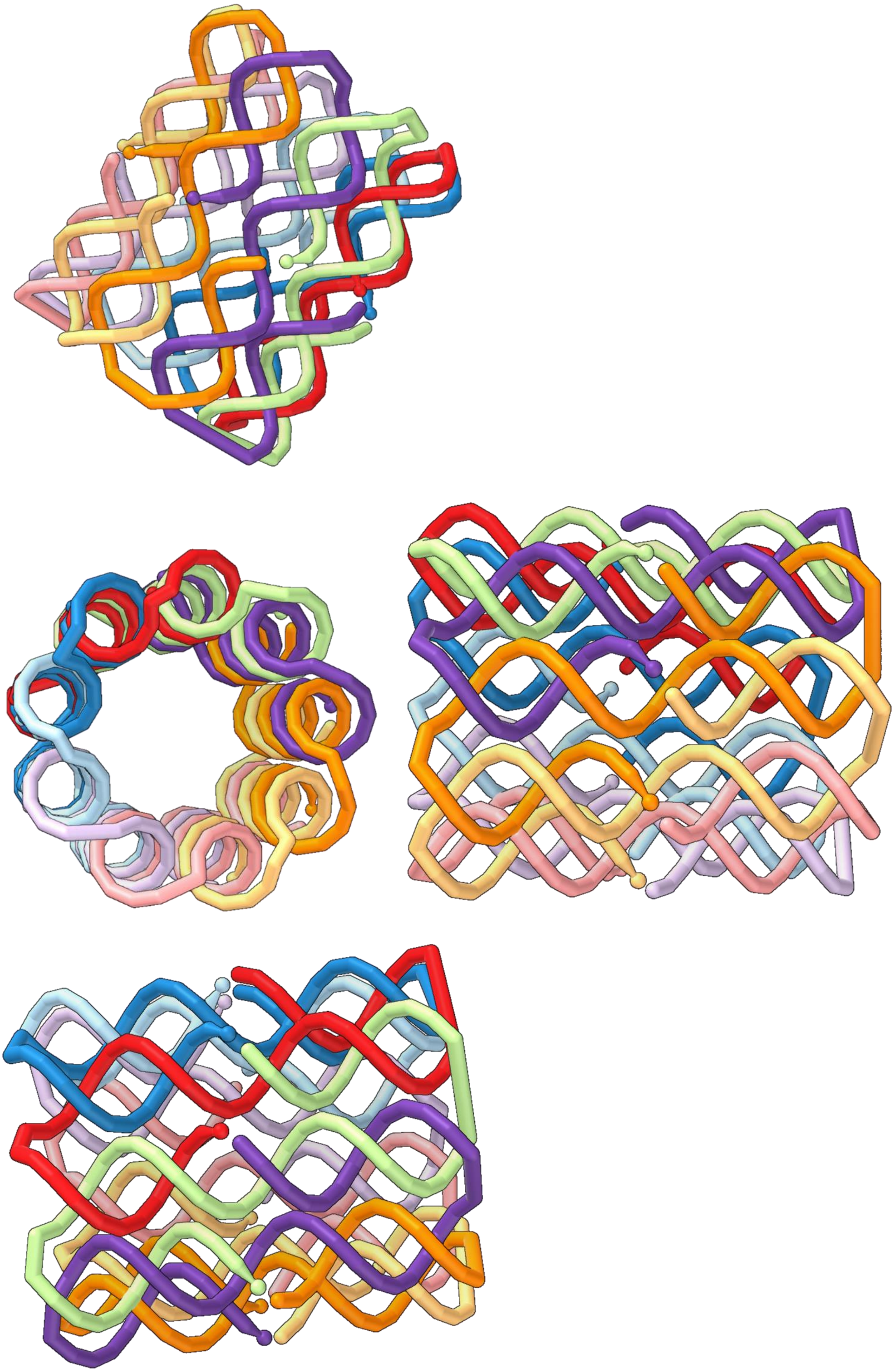

### Surface Plasmon Resonance (SPR) Sensor

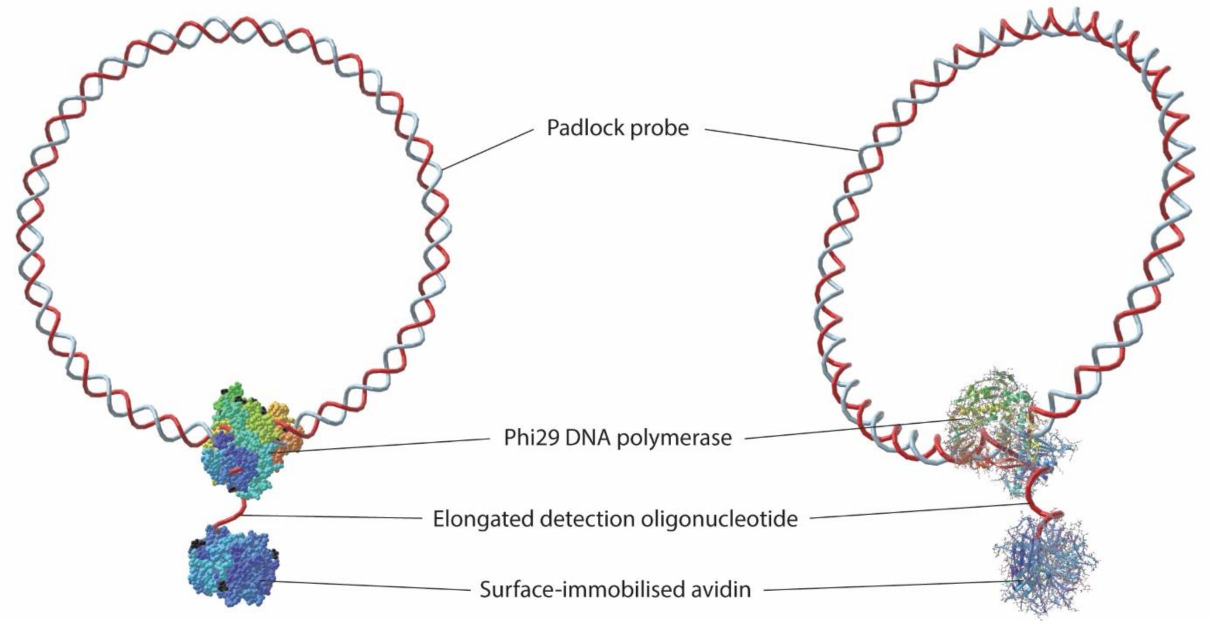


### DNA nanorobot

The nanorobot has been designed so it can switch between two conformations by attaching apta-switches in key parts of the design.

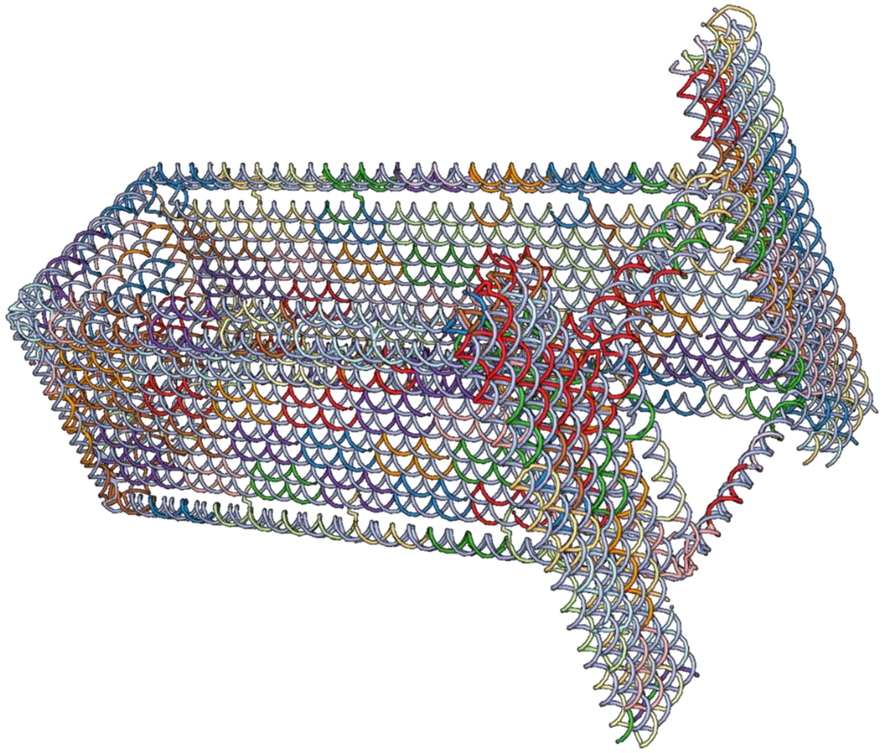

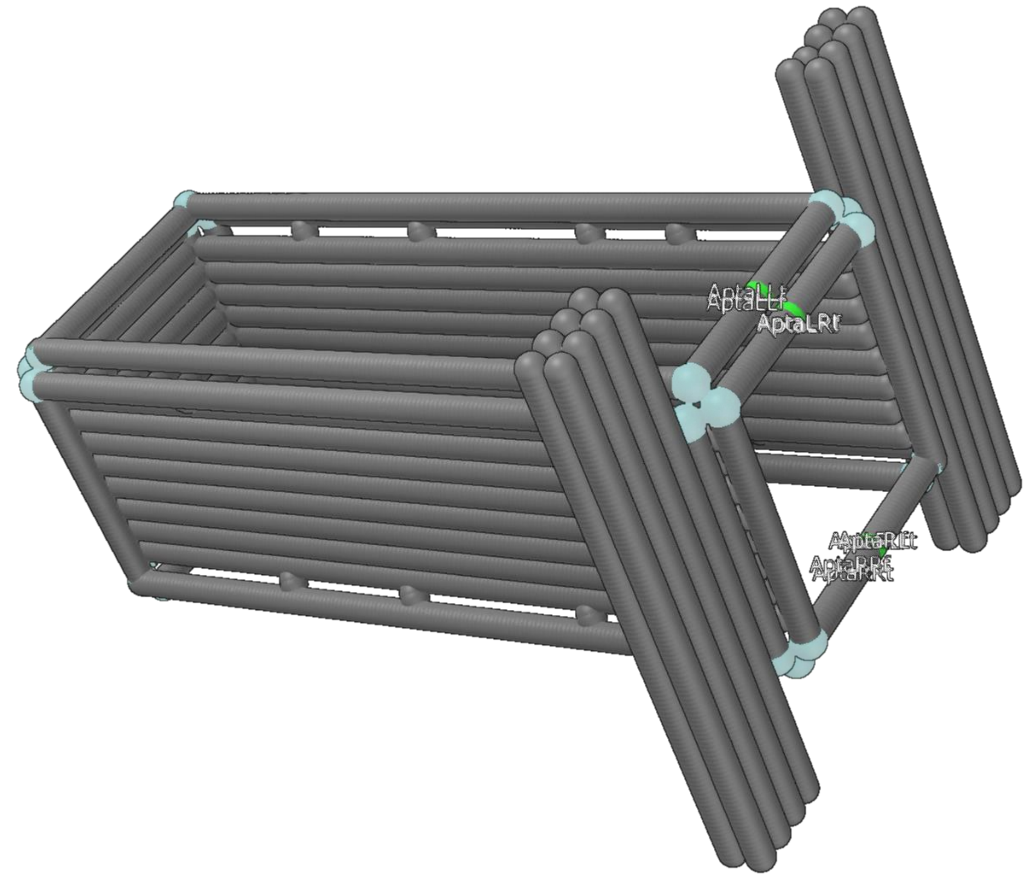


**Poly-T regions are represented as blue spheres. Nucleotides connecting with the apta-switches are represented in green and its label is depicted.**

### Design steps

1. Creation of a cuboid using the wireframe editor and a lattice that will cover a face

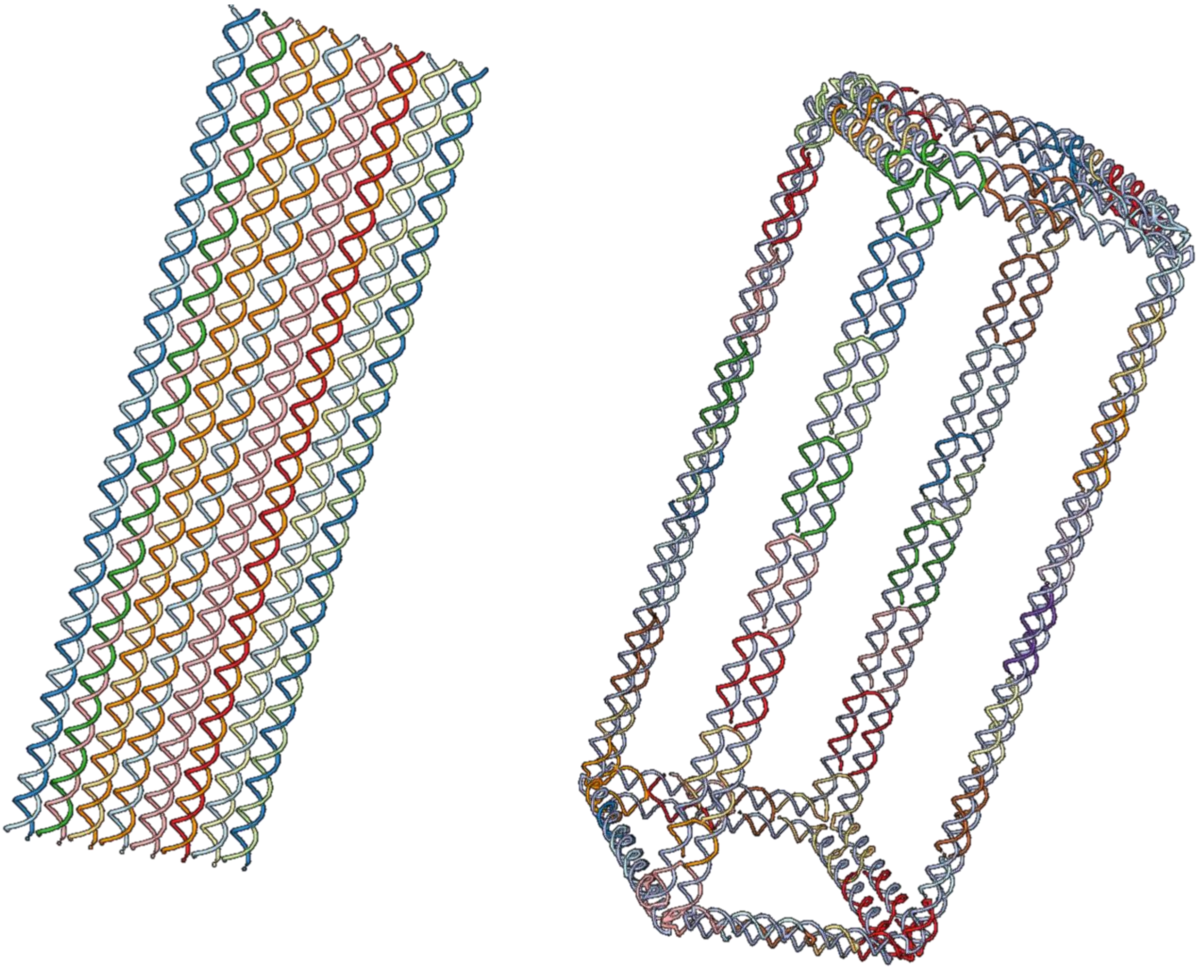

2. Lattice is placed to cover one of the faces. It can then be copied and place to cover another face

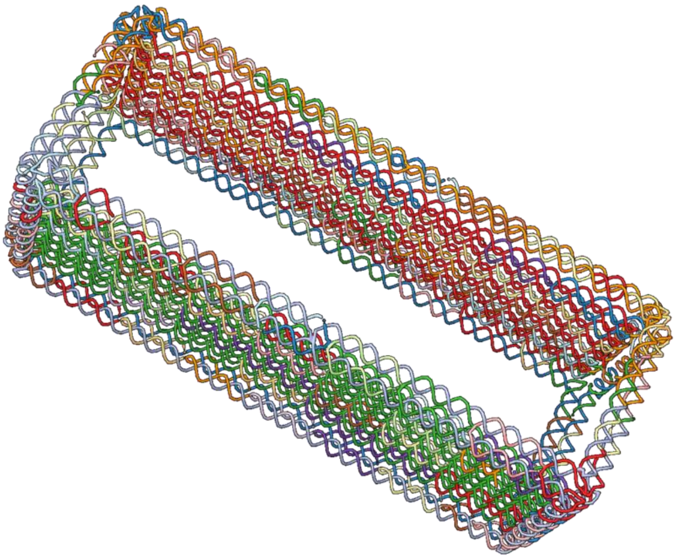

3. The scaffold is manually routed through the solid faces.

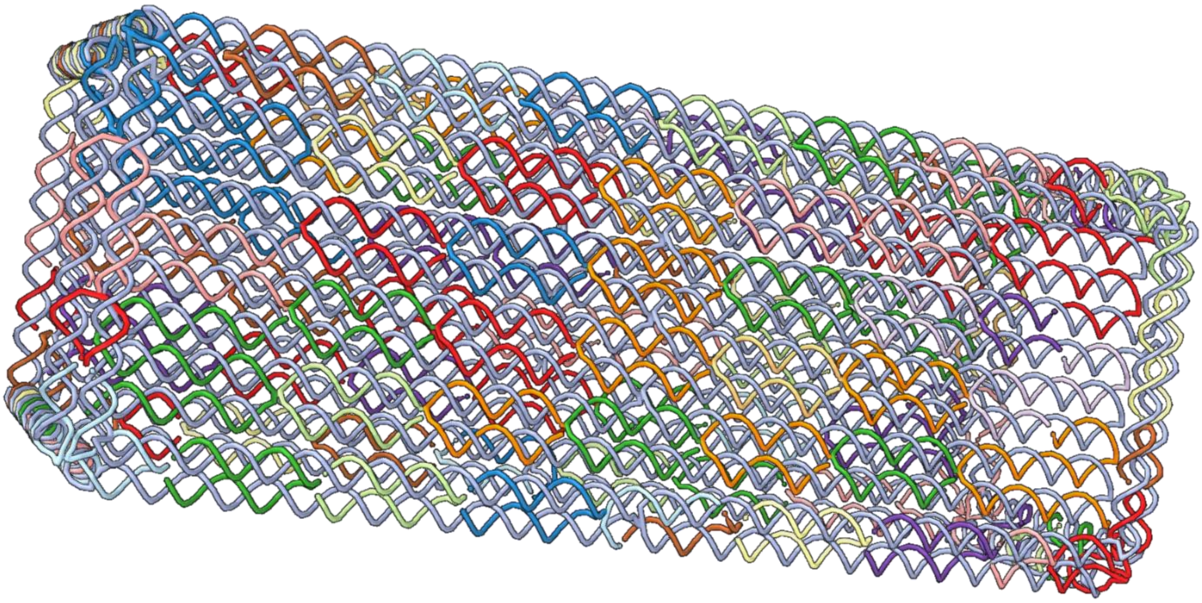

4. More DNA layers are added:

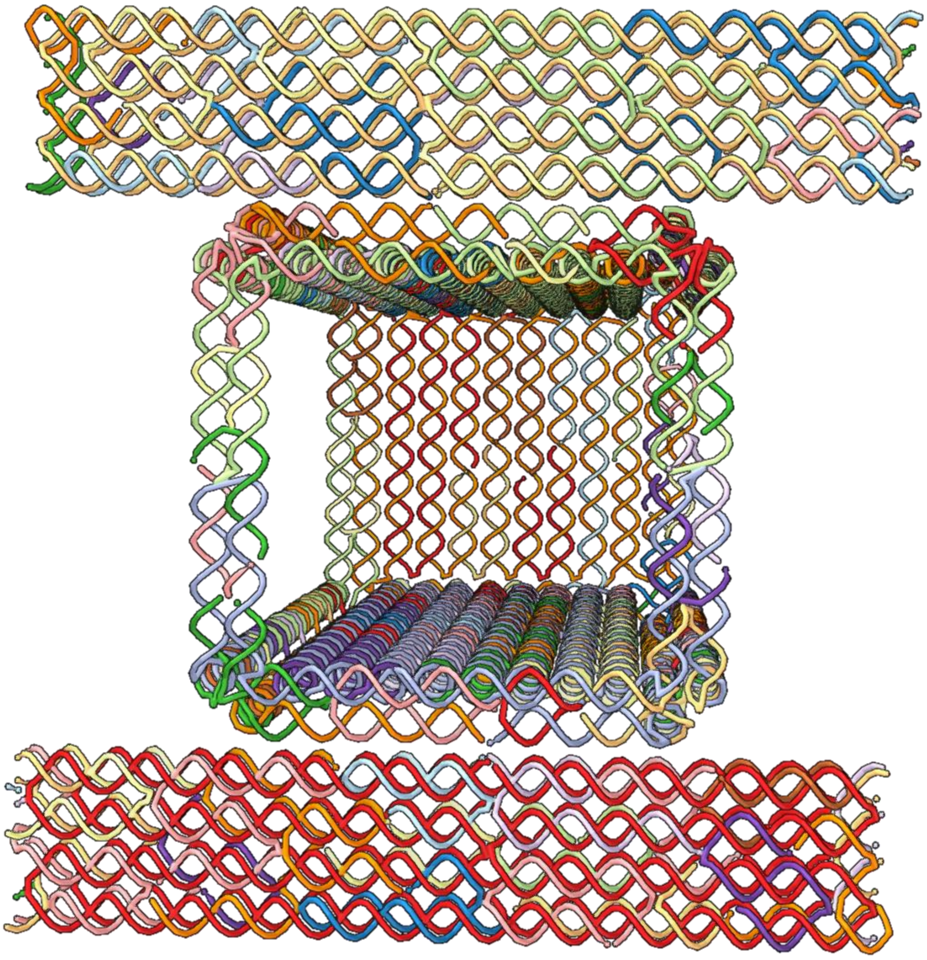

5. The scaffold (highlighted in green) is manually routed through the entire design by concatenating single-strands:

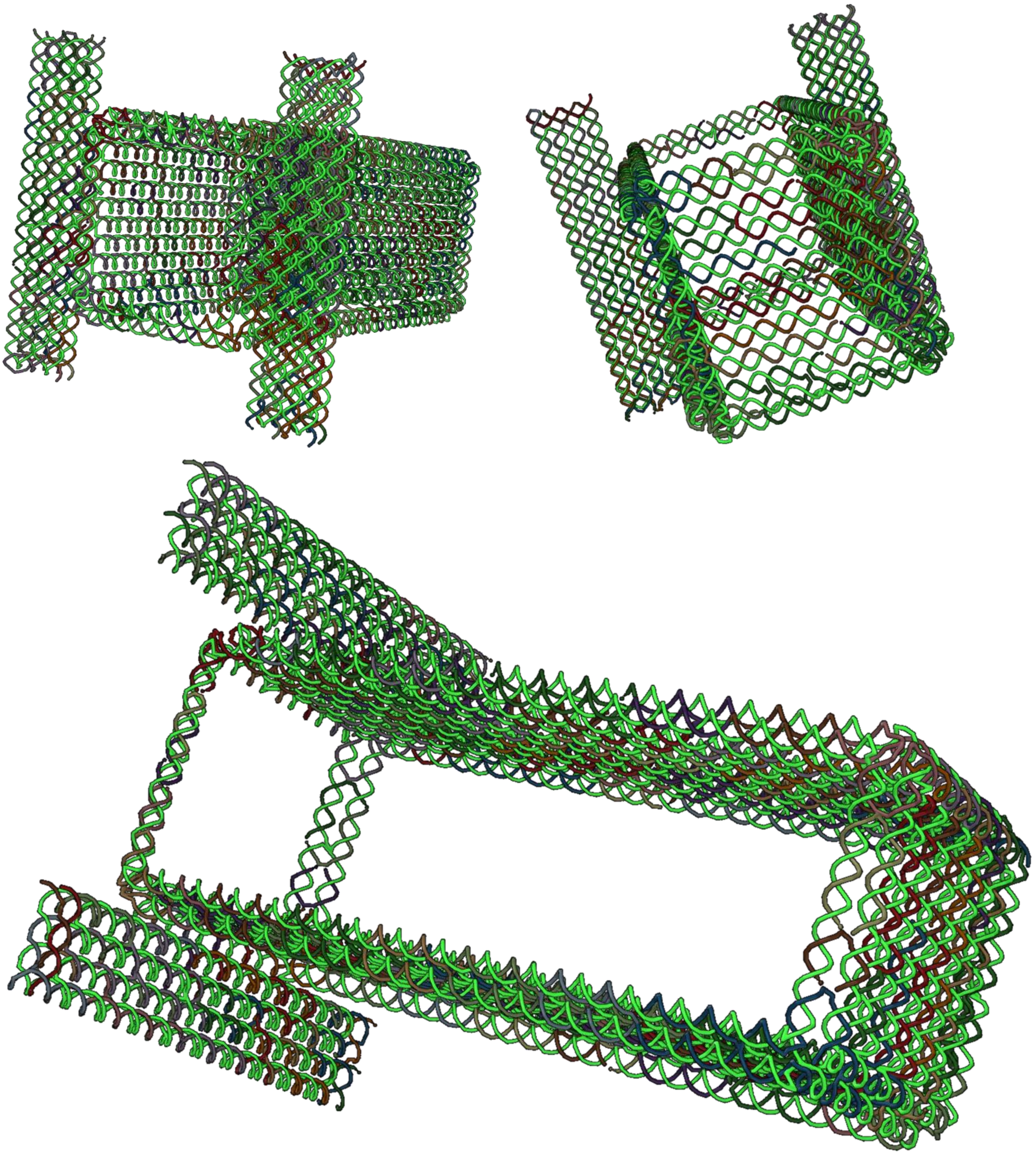

6. Visual inspection allows to determine where the apta-switches should be connected, and the corresponding nucleotides are tagged.

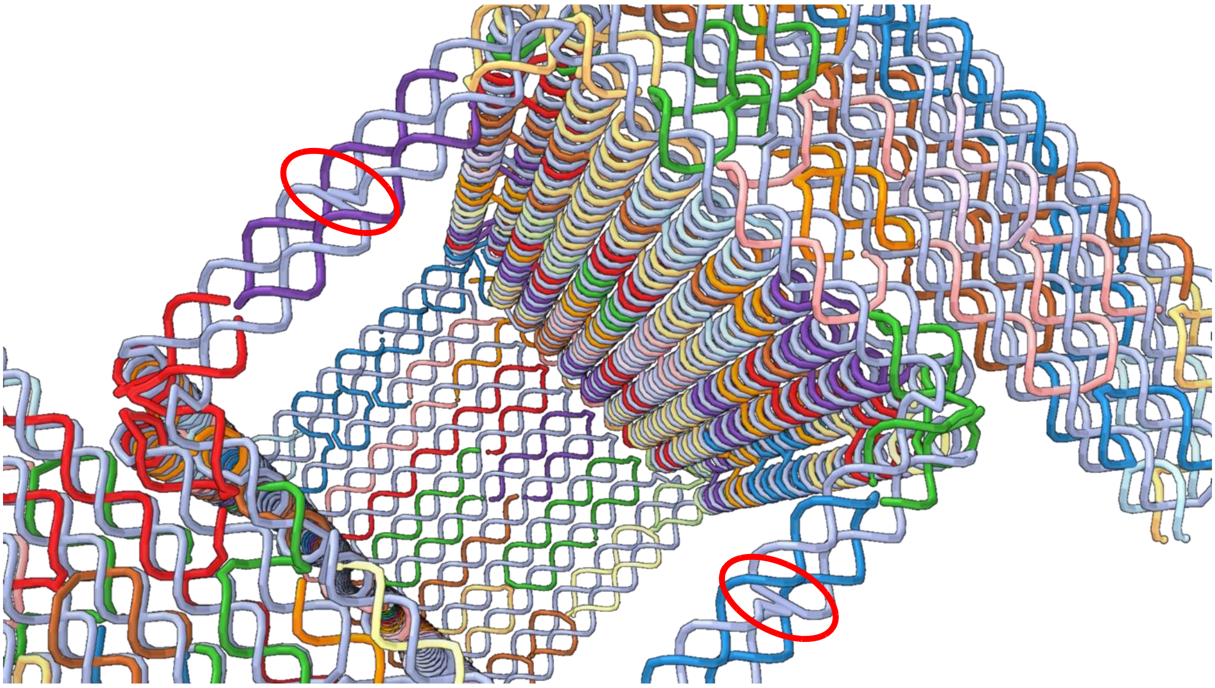

**Because the scaffold has crossovers at highlighted positions, breaking the staple strands there would result in the cuboid losing stability and opening-up.**

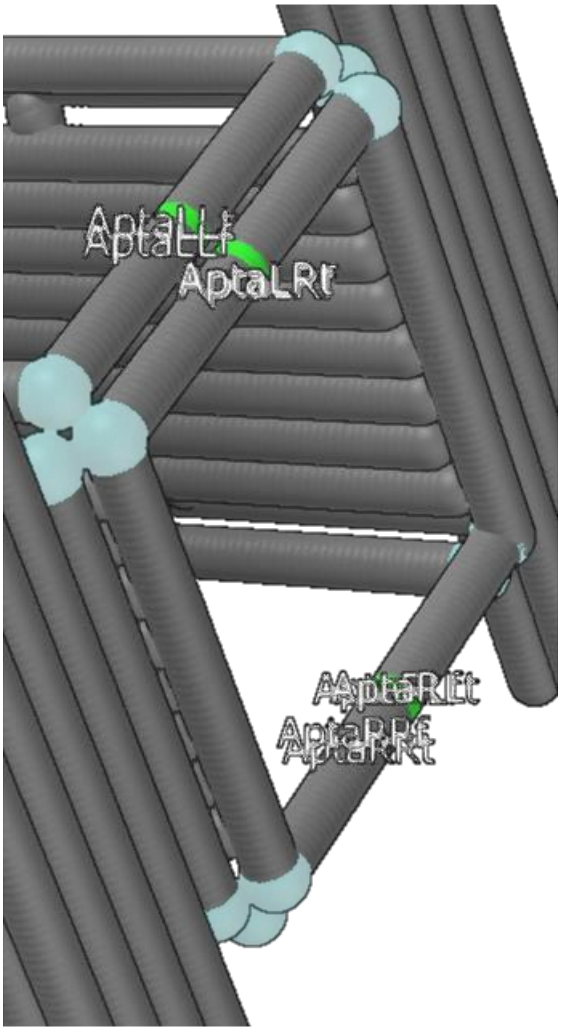


**Staples are modified and tagged for connection to apta-switches, to allow the cuboid to open up.**
