## Supplementary material for "Adenita: Interactive 3D modeling and visualization of DNA Nanostructures": User Documentation

### ADENITA: QUICK-START GUIDE

*This document is a quick-start guide to get you started designing DNA nanostructures on Adenita.*

#### REQUIREMENTS

Adenita is available for Windows and Linux. A dedicated graphics card is recommended.

#### FIRST STEPS

Adenita has been developed as a SAMSON plugin. You have to install **SAMSON 0.7.0** to use Adenita. To install SAMSON 0.7.0 please make sure you tick the box "Install an older version" when installing SAMSON.

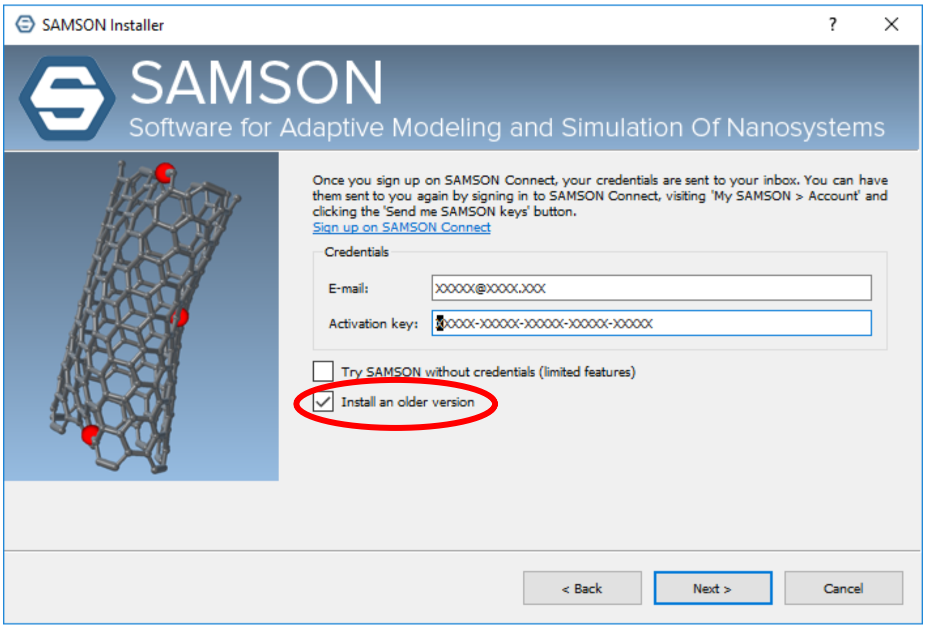


Afterwards you can select Adenita from a variety of plugins from the Samson's Element website. Adenita can be combined with any of them! If you are unfamiliar with SAMSON, check out their tutorials.

Adenita will work on **SAMSON 0.8.5**, but this branch is still being tested. This document is based on the 0.7.0 version.

#### VIDEO TUTORIALS

We have created a series of video tutorials where users can check some of Adenita's features and learn how to use it:

| | |
|---|---|
| Getting Started | http://bit.ly/35scneo |
| Creating ssDNA and dsDNA | http://bit.ly/2pKButL |
| Creating nanotubes and untwisting | http://bit.ly/35wF2PE |
| Creating square/honeycomb lattices and connecting with ssDNA | http://bit.ly/2qv6izb |
| Creating DNA wireframe structures with the editor and loading all atom models | http://bit.ly/2XKnzAy |
| Load and visualize Proteins (PDB) while working with DNA nanostructures | http://bit.ly/2OdUUk9 |
| Highlighting and tagging | http://bit.ly/2Odrxya |

#### FEATURES

##### CREATE DNA NANOSTRUCTURES

Use different editors to create dsDNA, nanotubes, lattices or wireframe nanostructures using the Daedalus algorithm.

##### SAVE COMPONENTS OF YOUR DESIGN, OR THE ENTIRE DESIGN

You can save the entire workspace using SAMSON's save function. If you want to save components of your design for later reuse use Adenita's saving function (*.adnpart*).

##### IMPORT A DNA NANOSTRUCTURE FROM CADNANO, OR LOAD A PREVIOUS DESIGN

Through the main UI you can load a Cadnano design, or component saved as *.adnpart*. You can combine as many components as your graphics card and CPU can afford.

##### EXPORT YOUR DESIGN

Users can also export your design as a list of sequences or in oxDNA format for simulations.

#### ADENITA UI

Once you have installed Adenita the main UI should be located with other applications:

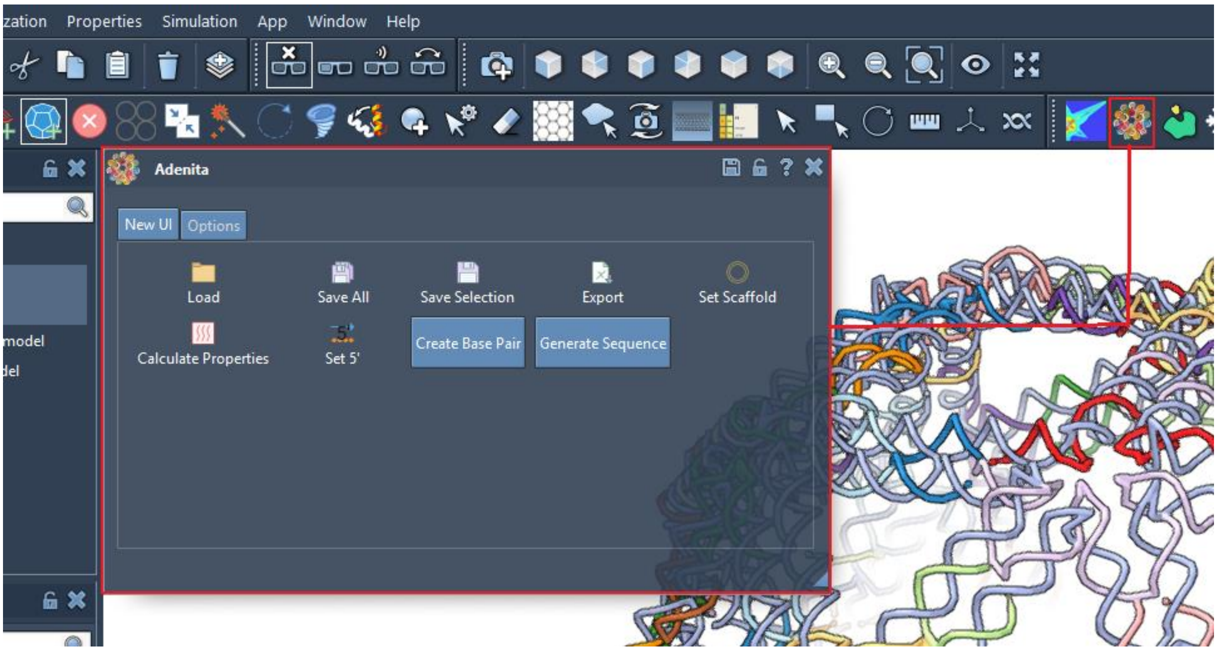


Different editors can be found on the editor menu:

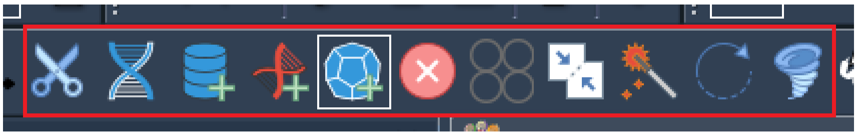

Users can also set some options:

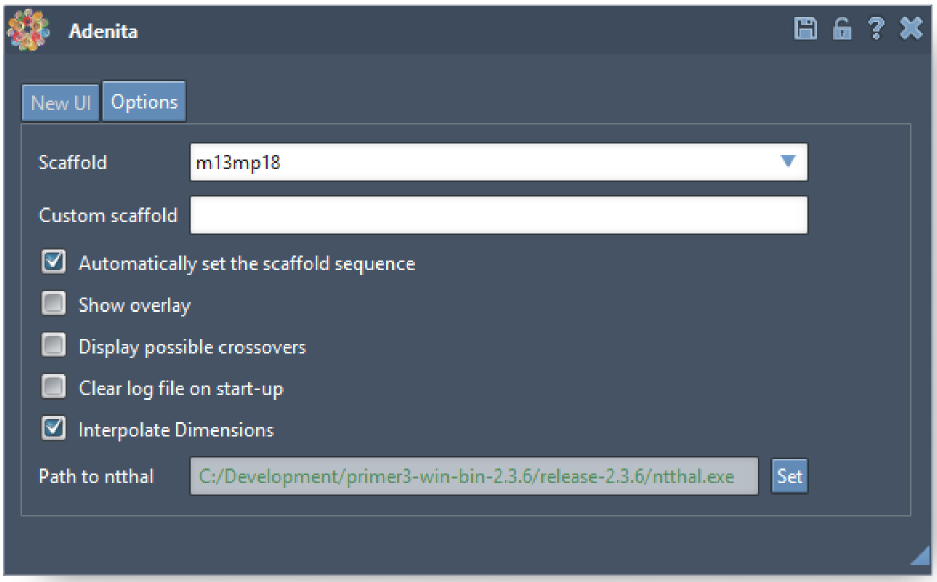


#### DESCRIPTION OF THE MAIN FUNCTIONS

##### MAIN UI

The following functions can be accessed through the main UI:

- 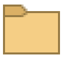 Load a DNA nanostructure from a file. Possible choices are a Cadnano design (for Cadnano 2.5) as *.json*, a mesh in *.ply* (will be loaded using the Daedalus algorithm), or a *.adnpart* or *.adn* (custom Adenita formats). This option allows users to load components into a workspace (loading using SAMSON files will create a new document).
- 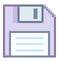 Users can chose to save a component for later use in our custom format (*.adnpart*).
- 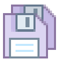 Users can save all current DNA nanostructures in a *.adn* file, systems not handled through Adenita won't be saved.
- 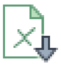 Options to export sequences as CSV file or in a format appropriate for oxDNA are available here.
- 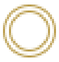 After selecting one or more components through SAMSON’s tree view, all scaffolds from the selection will be assigned a sequence specified through the 'Options' menu, scaffold nucleotide's pairs will also be assigned the complementary base. Strands can be marked as “scaffold” through SAMSON’s inspector.
- 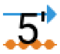

It is possible to set any nucleotide as the new 5' of its single strand.
- 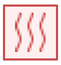 If a path to ntthal has been specified in the Options menu, it will be used to calculate the melting temperatures and Gibbs free energies of all binding regions of a selected component.

#### EDITORS

- Breaks the bond between two consecutive nucleotides of the same strand.
- Deletes nucleotides or base pairs, depending on the chosen visualization scale.
- Merges single or double strands. If strand ends are selected, they will be connected in the appropriate direction (5' to 3'). If nucleotides that are not 5' or 3' are selected, the strands will be broken to reconnect them at chosen points. It is also possible to insert a new double or single strand along the connection.
- Reorganize several components into one, or reassign single and double strands to other components. List of components and strands needs to be updated manually.
- Modifies the twist angle of a double-strand along the helical axis.
- Tag nucleotides or modify its base.
- Remove entire the twist of a double strand locally to observe the single strands that compose it as parallel lines.
- Add a new single or double strand as a component to the design. They can also be circular.
- Add a lattice of double strands as a component.
- Add a nanotube composed of double strands.
- Generate a wireframe from the given shapes and add it to the design (uses the [Daedalus](http://daedalus-dna-origami.org/) algorithm).

#### WHERE TO GET HELP

You can reach Adenita's developers at:

https://github.com/edellano/Adenita-SAMSON-Edition-Win- (Windows)

https://github.com/edellano/Adenita-SAMSON-Edition-Linux (Linux)

*Icons from: https://icons8.com/icons.*
